# The CPEB ortholog Orb2 regulates brain size through the TRIM-NHL RNA-binding protein, Brain tumor

**DOI:** 10.1101/2025.07.18.665534

**Authors:** Taylor Hailstock, Joseph Buehler, Beverly V. Robinson, Timothy C.H. Low, Yuqing Hua, Temitope H. Adebambo, Jack Govaerts, Todd A. Schoborg, Howard D. Lipshitz, Dorothy A. Lerit

## Abstract

Neurodevelopment requires precise translational control, the disruption of which is implicated in various neurological disorders, including developmental delays, intellectual disability, and microcephaly. We report a novel role for the *Drosophila* CPEB-family protein Orb2, a translational regulator, in controlling brain size in a dose dependent manner. Loss of *orb2* results in larval brain hypotrophy, whereas *orb2* overexpression causes brain overgrowth. We demonstrate that *orb2* is required for neural stem cell development from embryonic stages through larval neurogenesis. Structure-function analysis reveals that Orb2 RNA-binding activity promotes brain growth, while its poly-Q and ZZ domains act to restrain overgrowth. Further genetic and biochemical evidence indicates that *orb2* functions upstream of the translational repressor Brain tumor (Brat), modulating Brat protein levels and, consequently, influencing brain size. These findings support a model wherein the antagonistic activities of Orb2 and Brat are critical for balanced brain growth during *Drosophila* neurodevelopment.

## INTRODUCTION

RNA-binding proteins (RBPs) regulate neurodevelopment by controlling neural stem cell (NSC) proliferation, differentiation, and maintenance through post-transcriptional gene regulation (Darnell, 2013; Lennox et al., 2018; Prashad & Gopal, 2021). RBPs are critical for regulating various aspects of stem cell function and homeostasis by directly modulating RNA metabolism, localization, splicing, and transcript turnover, which are essential for proper neuronal development and function (Lennox et al., 2018; Parra & Johnston, 2022). Consequently, mutations in RBPs are associated with intellectual disability, developmental delays, seizures, microcephaly, and other neurodevelopmental disorders (Allanson et al., 2009; Barkovich et al., 2012; Pirozzi et al., 2018).

The conserved cytoplasmic polyadenylation element-binding (CPEB) proteins are RBPs that orchestrate the translation of target transcripts through canonical CPE sites in the 3’ untranslated region (3’UTR; (Hake & Richter, 1994). While CPEB proteins, such as CPEB4, are implicated in neurodevelopmental disorders, such as autism spectrum disorder and Fragile X Syndrome (Udagawa et al., 2013; Garcia-Cabau et al., 2025), the complete repertoire of CPEB protein interactions and RNA targets, particularly across different tissues and developmental stages, remains incompletely understood.

*Drosophila* encodes two CPEB proteins from unique genes, Oo18 RNA-binding protein (Orb) and Orb2 (Lantz et al., 1992; Hafer et al., 2011). *Drosophila orb2* is orthologous to mammalian *CPEB2–4* and has known roles in mRNA localization and translation (Huang et al., 2006; Keleman et al., 2007; Hafer et al., 2011; Xu et al., 2014). Orb2 undergoes conformational changes to switch between a translational repressor and an activator dependent upon prion-like oligomerization, during which the N-terminal glutamine-rich (poly-Q) domain promotes self-association (Hake & Richter, 1994; Keleman et al., 2007; Si et al., 2010; Khan et al., 2015; Hervás et al., 2016). CPEB proteins like Orb2 bind CPE motifs in target mRNAs through RNA recognition motifs (RRMs; (Hake et al., 1998; Krüttner et al., 2012; Majumdar et al., 2012; Khan et al., 2015; Stepien et al., 2016). While the C-terminal ZZ-type zinc-finger domain (ZZ-domain) was previously implicated in RNA binding by anchoring CPEB to target transcripts (Hake et al., 1998), evidence also supports an alternative function in protein-protein binding (Legge et al., 2004; Merkel et al., 2013; Oroz et al., 2020). Consistently, recent data support a role for the ZZ domain in the recruitment of translational repressors to its target transcripts (Low et al., 2025).

CPEB proteins like Orb2 influence translation levels by regulating the polyadenylation status of target RNAs. Monomeric Orb2 binds to the deadenylase CG13928 to repress translation, while synaptic activity triggers Orb2 oligomerization and binding to CG4612, a predicted poly(A)-binding protein, to promote polyadenylation and translational activation of target mRNAs (Si et al., 2003a; Si et al., 2003b; Si et al., 2010; Majumdar et al., 2012; Khan et al., 2015). For example, recent work indicates that Orb2 promotes rare-codon mRNA translation in the brain (Stewart et al., 2024).

Cell type- and stage-specific expression of *orb2* enables dynamic mRNA stability and translation control required for neuronal function. Orb2 also autoregulates its own translation by binding CPE motifs in its 3’ UTR (Majumdar et al., 2012; Stepien et al., 2016). Despite dynamic changes in *orb2* expression and localization throughout neurogenesis (Hafer et al., 2011), Orb2 is best known for its roles in memory and learning. Orb2 has been extensively studied as a key synaptic regulator of long-term memory, wherein activity-induced aggregation facilitates sustained neuronal responses to learning. This molecular process is essential for stabilizing memory traces and supporting lasting behavioral adaptations (Si et al., 2003a; Keleman et al., 2007; Krüttner et al., 2012; Majumdar et al., 2012; Khan et al., 2015; Hervás et al., 2016; Li et al., 2016; Oroz et al., 2020; Kozlov et al., 2021; Kozlov et al., 2023). By comparison, relatively little is known about Orb2 functions during neurodevelopment.

*Drosophila* neurodevelopment commences during embryogenesis, when NSCs are specified by spontaneous delamination from the neuroepithelium (Hartenstein & Campos-Ortega, 1984; Doe, 1992). After delamination, NSCs undergo repeated rounds of asymmetric cell division to generate a self-renewing stem cell and a ganglion mother cell fated for differentiation into neuronal or glial subtypes (Broadus & Doe, 1997; Cabernard & Doe, 2009). The balance between self-renewal versus differentiation is governed by precise segregation of cell fate determinants and is vital to maintain homeostasis in neurogenesis, where deregulation can lead to tumorigenesis, neurodegeneration, or disorders such as microcephaly (Wang et al., 2016).

NSCs enter quiescence in late embryonic stages and resume proliferation during the first larval instar, termed NSC reactivation (Truman & Bate, 1988; Maurange & Gould, 2005). While about 10% of the neurons in the adult brain are born during embryonic stages, the remaining 90% of neurons arise throughout larval development (Prokop & Technau, 1991). Defects in NSC specification, asymmetric division, or maintenance during early stages are associated with neurodevelopmental deficits (Gonzalez et al., 1990; Rujano et al., 2013; Gogendeau et al., 2015; Schoborg et al., 2015; Ramdas Nair et al., 2016; Vargas-Hurtado et al., 2019; Mannino et al., 2023). For example, loss of the translational repressor and tumor suppressor *brain tumor* (*brat*) leads to overexpansion of Type II NSCs, resulting in brain hypertrophy (Bello et al., 2006; Lee et al., 2006).

In this study, we examined the role of Orb2 in neurodevelopment and identified an unexpected requirement for Orb2 in the regulation of larval brain size. We find that Orb2 governs brain size in a dose-dependent manner reliant on its RNA-binding activity, with mutants showing smaller brains consistent with microcephaly. In contrast, *orb2* overexpression results in enlarged brains. We identify cell autonomous requirements for Orb2 to regulate brain size through the specification and maintenance of NSCs during embryonic and larval development. We further uncover that *orb2* functions upstream of *brat*, demonstrating that the opposing activities of these RBPs support balanced brain growth.

## RESULTS

### Orb2 is a homeostatic regulator of brain size

To investigate Orb2 contributions to neurodevelopment, we first quantified brain volumes from age-matched third instar larvae (Link et al., 2019; Hailstock et al., 2023). Strikingly, *orb2* mutant brains were ∼70% smaller than wild-type (WT) brains (Fig. 1A,B and D; *p*<0.0001 by one-way ANOVA). These data indicate Orb2 is required for brain growth. To test if *orb2* overexpression was sufficient to enlarge larval brains, we drove expression of a *UAS-orb2-GFP* transgene (Krüttner et al., 2012) using *actin-GAL4* (*act*-*GAL4*; hereafter, *orb2^OE^*) and confirmed overexpression via western blot (Fig. S1A,B; *p*<0.01 by two-tailed t-test). Indeed, *orb2^OE^*brains showed an overgrowth phenotype with two-fold increased volumes relative to WT (Fig. 1C,D; *p*<0.0001 by one-way ANOVA). We conclude that Orb2 regulates larval brain size in a dose-dependent manner.

**Figure 1.**
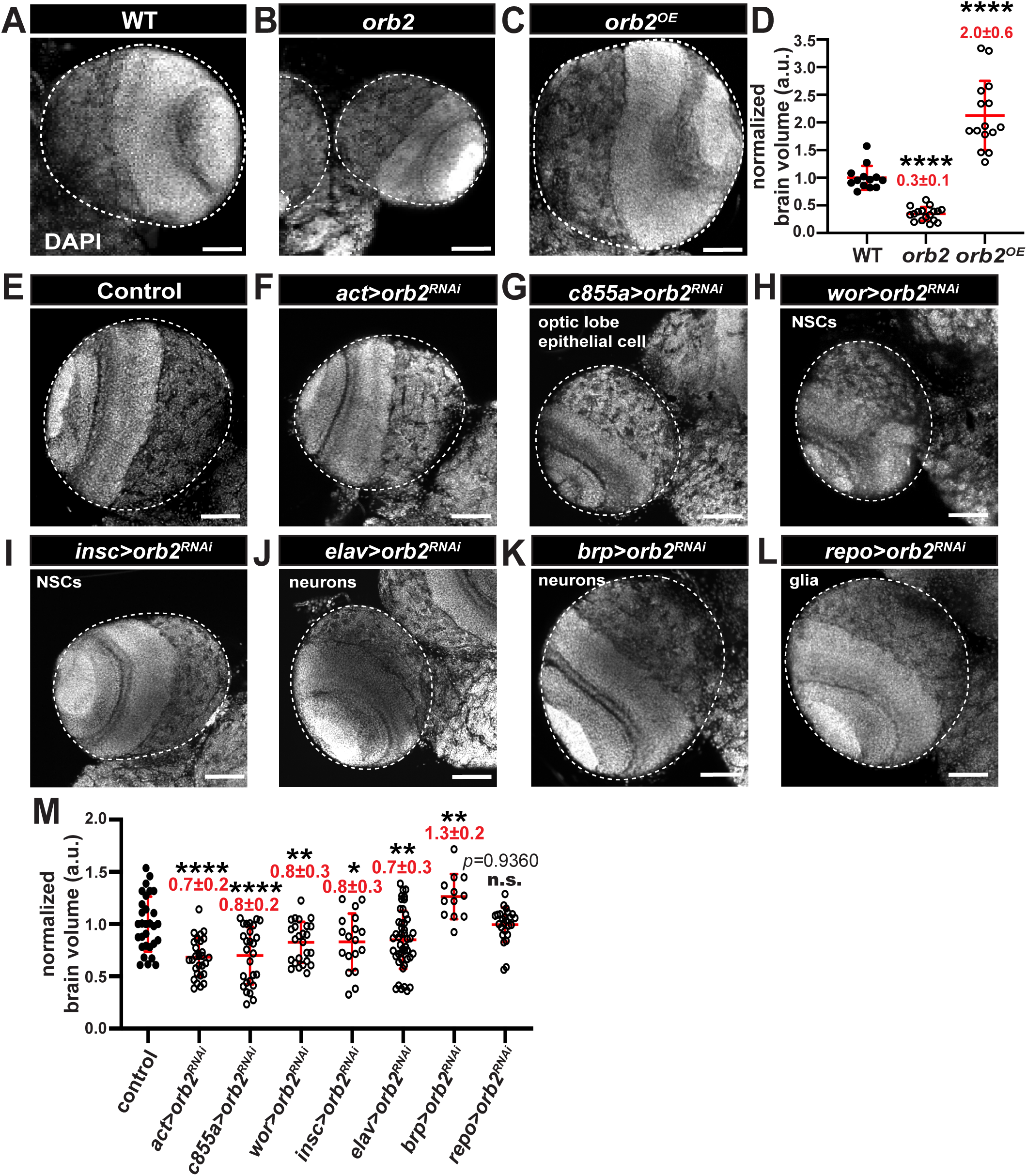
***orb2* is necessary and sufficient for brain growth**. Images show the optic lobe region (dashed circles) from age-matched third instar larval brains of the indicated genotypes stained for DAPI. (A) WT, (B) *orb2* null (*orb2^Δ36^*) mutants show reduced brain size, and (C) *act-GAL4* driven expression of *UAS-orb2* (*orb2^OE^*) results in increased brain volumes, as quantified in (D). Graph shows normalized brain volumes, where each dot represents a measurement of a single optic lobe from N=13 WT, 17 *orb2*, and 15 *orb2^OE^* brains. Data shown are pooled from 2 (WT, *orb2^OE^*) or 3 (*orb2*) independent biological replicates. Representative images of brains expressing (E) *UAS-orb2^RNAi^* (no-GAL4 control) alone or (F–L) in combination with the specified GAL4. Reduced brain volume was observed following depletion of *orb2* by (F) ubiquitous expression via *act-GAL4*, (G) within neuroepithelia cells of the outer proliferation center via *c855a*-*GAL4*, in neural stem cells via (H) *wor-GAL4* or (I) *insc-*GAL4, and in (J) and neurons via *elav-GAL4*. In contrast, *orb2* depletion in (K) mature neurons via *brp-GAL4* increased brain size, while (L) brain volume was unaffected in glia using *repo*-GAL4. (M) Quantification of brain volume normalized to controls, where each dot represents a single measurement from a single optic lobe from N= 30 control, 20 *act>orb2^RNAi^*, 25 *wor> orb2^RNAi^*, 18 *insc>orb2^RNAi^*, 47 *elav>orb2^RNAi^*, 27 *c855a>orb2^RNAi^*, 12 *brp>orb2^RNAi^*, and 25 *repo> orb2^RNAi^* pooled from 2 (control, *wor*, *insc*, *c855a*, and *repo*) or 3 (*act*, *elav*, *brp*) independent biological replicates. Significance was determined by one-way ANOVA. n.s., not significant; \**p*<0.05; \*\**p*<0.01; \*\*\**p*<0.001; \*\*\*\**p*<0.0001. Scale bars: 50 μm.

### Orb2 is required in neuronal lineages to support brain size

The larval brain comprises distinct cellular identities, including NSCs, marked by Miranda (Mira, (Shen et al., 1997)) or Inscuteable (Insc, (Kraut et al., 1996)); glia, marked by the *reversed polarity* (*repo*) gene (Xiong et al., 1994); and neurons, marked by *embryonic lethal abnormal vision* (*elav*) (Robinow & White, 1988). To characterize the cellular etiology of *orb2*-dependent microcephaly, we selectively depleted *orb2* in defined cellular lineages within the larval brain, including NSCs, neurons, glia, and the neuroepithelium using shRNA (*orb2^RNAi^*) and a panel of GAL4 drivers (Hrdlicka et al., 2002). We then compared brain volumes from age-matched larval brains to controls that carry the RNAi transgene, but not a GAL4 driver; hereafter, ‘no GAL4’ controls.

We first confirmed that uniform expression of *orb2^RNAi^*with *act-GAL4* was sufficient to reduce brain volume (Fig. 1E,F, and M). Depletion of *orb2* by RNAi reduced brain volume to a lesser degree than the *orb2* null mutation, likely due to the partial depletion of *orb2* activity, as shown by immunoblot (∼50% reduced; Fig. S1C,D). Similar reductions in brain volume were also detected when *orb2* was depleted in neurogenic lineages: neuroepithelia cells, which give rise to medulla neuroblasts of the optic lobe (c*855a-GAL4* (Egger et al., 2007; Wang et al., 2011; Shard et al., 2020) *p*<0.0001), NSCs (*wor*-*GAL4* (Ashraf et al., 1999); *p*<0.05 or *insc-GAL4*; p<0.01), or neurons (e*lav-GAL*4; *p*<0.01), underscoring requirements for *orb2* in these lineages to promote brain growth (Fig.1G–J and M; by one-way ANOVA). In contrast, no difference in brain volume was noted upon depletion of *orb2* in glia via *repo-GAL4* (Fig. 1L, M; n.s. by one-way ANOVA). These data indicate that Orb2 facilitates the establishment of normal brain size in neural progenitors and their neuronal progeny.

### *orb2* expression becomes progressively restricted to mature neurons

We next examined where Orb2 is expressed during larval neurodevelopment. To assess Orb2 distributions in live larval brains, we imaged third instar larval brains expressing endogenously-tagged *Orb2-GFP*, previously shown to functionally restore *orb2* activity (Krüttner et al., 2012), relative to an mCherry nuclear reporter driven by cell type specific drivers. Through this approach, we found that Orb2 is not detectable within glia, marked by *NP2222-GAL4* or *repo-GAL4*, nor in the optic lobe neuroepithelium or lamina precursor cells, marked by *c855a-GAL4* (Fig. 2A). In contrast, *Orb2-GFP* was coincident within a subset of smaller *insc-GAL4* expressing cells (*arrows*; Fig. 2A and discussed below). However, at this stage, Orb2 was not detected within third instar larval NSCs, characterized by their distinctive, large nuclei (*dashed circles*; Fig. 2A). Taken together, these data show that *Orb2-GFP* positive cells are non-glial descendants of the NSC lineage (i.e., neurons).

**Figure 2.**
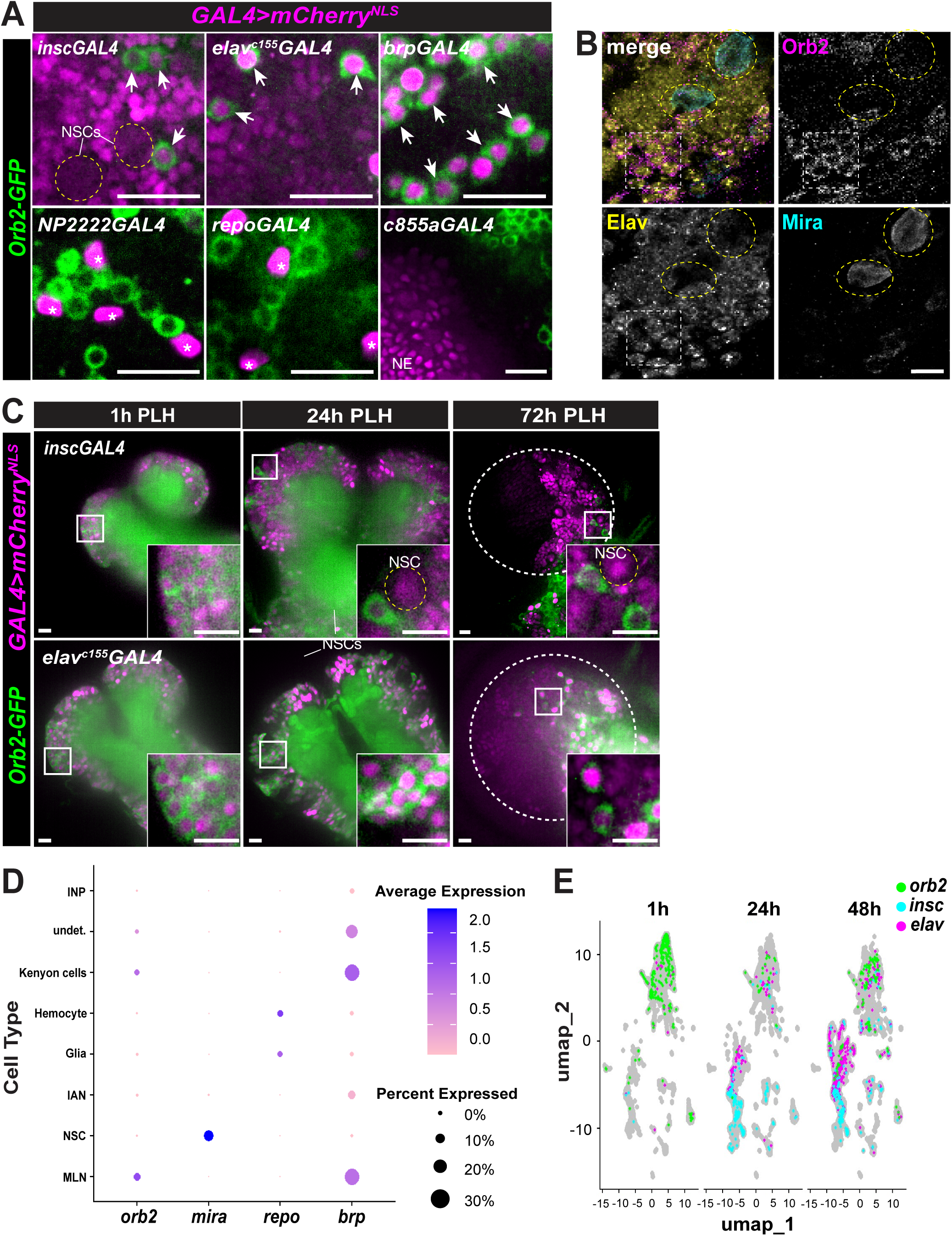
Orb2 becomes restricted to mature neurons. (A) Live images of endogenous *Orb2-GFP* localization relative to a *UAS*-*mCherry_NLS_* reporter (magenta) driven by the indicated GAL4 lines: NSCs (*insc*), neurons (*elav^c155^*), mature neurons (*brp*), glia (*NP2222* and *repo*), and neuroepithlelial cells (*c855a*). Dashed circles mark NSCs, asterisks mark glia, and arrows highlight cells positive for both Orb2 and the mCherry reporter; NE, neuroepithelium. (B) Representative immunofluorescence images of third larval instar brains stained for Orb2 (magenta) localization relative to neurons (Elav, yellow) and NSCs (Mira, cyan). Boxed area marks Elav and Orb2-positive neurons. (C) Live images of Orb2 (green) distributions relative to NSCs (*insc*) or neurons (*elav;* magenta) throughout the first (1h), second (24h), and third (72h) larval instars; time is hours post larval hatching (PLH). Orb2 exclusion from NSCs (yellow circles) becomes apparent at 24h and 72h PLH. Over time, Orb2 enriches to a subset of *elav+* neurons (72h). (D) Seraut dot plot comparing the average expression of the indicated genes (x-axis) across all larval stages relative to known cell type biomarkers (y-axis). (E) UMAP plot of scRNA-seq data (Corrales et al., 2022) comparing cells from the first (1h), second (24h), and third (72h) instar larval brains with normalized expression values of *orb2* >3.0 (green), *insc* >0.75 (cyan), and *elav* >3.5 (magenta). Scale bars: 10 µm.

Prior work indicates that Orb2 is primarily localized to the cytoplasm of *Drosophila* neurons, including mature glutamatergic neurons (Si et al., 2003a; Keleman et al., 2007; Li et al., 2016; Stewart et al., 2024). Consistently, *Orb2-GFP* expression was detected within a subgroup of *elav^c155^-*positive cells determined to be mature larval neurons given strong co-expression with *bruchpilot* (*brp*)-GAL4 (Wagh et al., 2006) (*arrows*; Fig. 2A). Immunostaining confirmed that endogenous Orb2, as detected with anti-Orb2 antibodies, labels a subset of Elav+ neurons, but not NSCs, within third instar larval brains (*box*; Fig. 2B).

Because loss of *orb2* activity in NSCs results in impaired brain growth (Figure 1), we next assayed whether Orb2 enriches within NSCs at earlier stages of development. Low levels of Orb2 were detected within embryonic Mira+ dorsoanterior neural progenitors, consistent with a possible early function in neural lineages (Fig. S1E). Upon examining first, second, and third instar stages, we noted that the distribution of Orb2 becomes progressively more restricted over time (Fig. 2C). At the first larval instar (1 hr post-larval hatching, PLH), *Orb2-GFP* was detected within subsets of *insc-* or *elav*-expressing cells. However, the exclusion of *Orb2-GFP* from NSCs became apparent in second (24h PLH) and third (72h PLH) instar larval brains (*dashed circles*; Figure 2C).

To further examine *orb2* expression, we analyzed published single cell transcriptomics (scRNA-seq) data of the *Drosophila* larval central nervous system across the first, second, and third instar stages (Corrales et al., 2022). In accordance with our protein reporter assay, *orb2* RNA was found to be expressed within *brp*+ mature larval neurons (MLN), including Kenyon cells of the mushroom body. In contrast, levels of *orb2* RNA within *mira*+ NSCs were lower (Figure 2D). Examination of UMAP clustering patterns likewise indicated that *orb2* mRNA distributions are more similar to *elav* than the NSC marker *insc* across larval development (Figure 2E). Thus, during larval development, *orb2* and *elav* are expressed within cells with similar transcriptional profiles.

Together, these data suggest that *orb2* expression becomes increasingly restricted within neurons as development ensues. To test if *orb2* is necessary in mature neurons to promote brain growth, we drove *orb2^RNAi^* expression using a *brp*-*GAL4.* Unexpectedly, depletion of *orb2* in mature neurons was not sufficient to reduce brain size, but caused overgrowth (Fig. 1K,M; \*\**p*<0.01 by one-way ANOVA). In contrast to *brp*, *elav* expresses strongly not only in mature neurons, but also in NSCs and their immediate progeny (O’Neill & Rusan, 2022). We conclude that *orb2* is required in neurogenic lineages to regulate brain size. Our data further suggest that Orb2 likely functions within neural progenitors and newly born neurons to promote brain growth. These findings underscore a previously undefined role for Orb2 in neurodevelopment.

### Multiple Orb2 domains instruct brain size

*orb2* encodes two protein isoforms, Orb2A (predicted molecular weight, ∼60kDa) and Orb2B (∼75kDa), which differ only in the length of an unstructured N-terminal region with unknown function (Hafer et al., 2011). As previously noted, Orb2B is the predominant isoform in third instar larval brains (Fig. S1C; (Mastushita-Sakai et al., 2010; Hafer et al., 2011; Majumdar et al., 2012)). Because multiple domains within Orb2 support its function as an RBP implicated in translational regulation, we performed structure-function analysis to identify which domain(s) regulate larval brain size (Fig. 3A; (Krüttner et al., 2012; Majumdar et al., 2012; Khan et al., 2015; Stepien et al., 2016).

**Figure 3.**
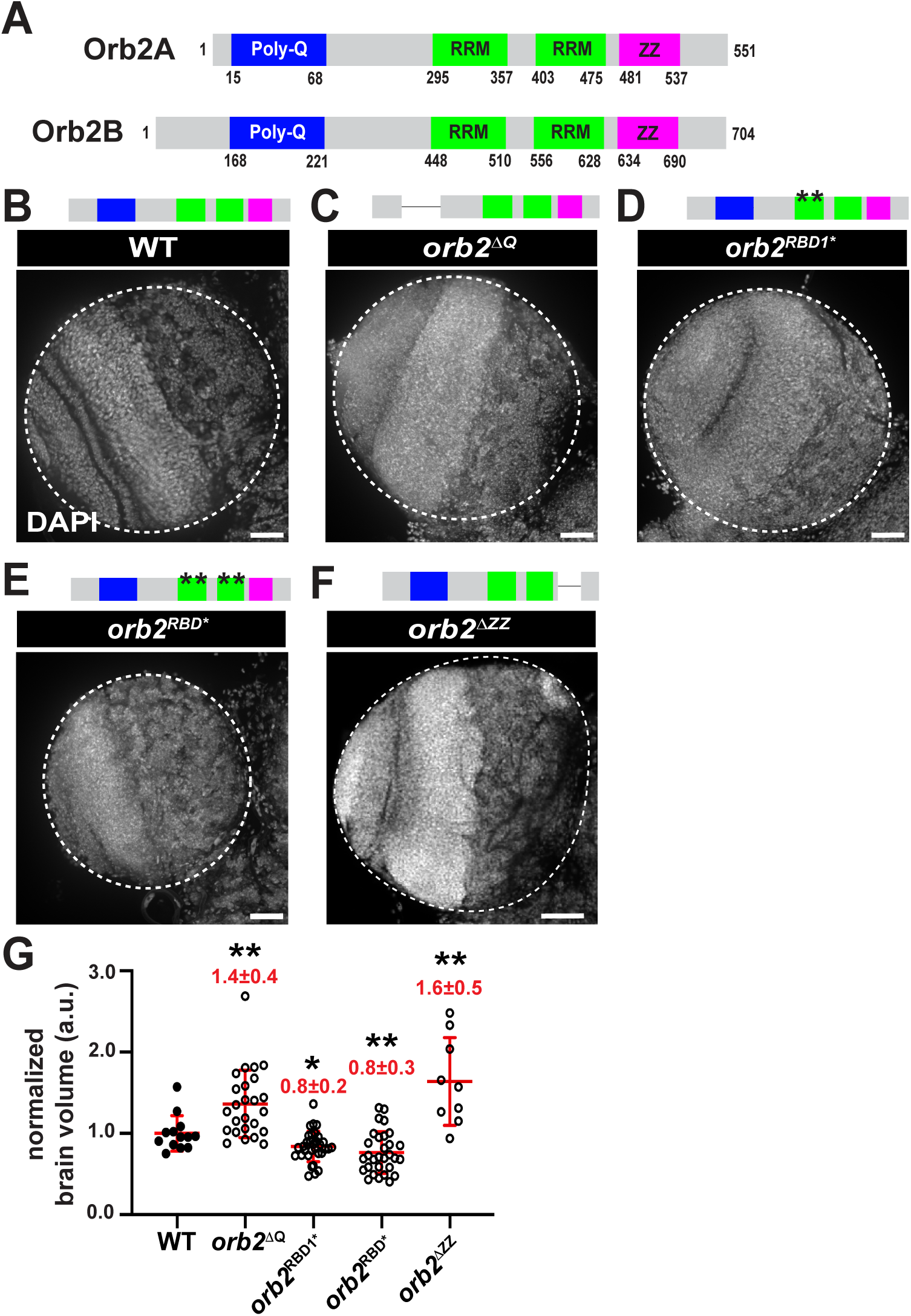
Multiple Orb2 domains instruct brain size. (A) Schematic of annotated Orb2 domains: poly-Q domain (glutamine-rich, blue), RNA-recognition motif (RRM, green), and ZZ domain (magenta). (B–G) Representative images of age-matched third larval instar brains of the indicated genotypes stained for DAPI. A schematic of Orb2 structure is shown above each image (astericks denote single AA mutations). (H) Quantification of brain volumes, where each dot represents a single measurement from a single optic lobe from N=13 WT, 25 *orb2^ΔQ^*, 32 *orb2^RBD1*^*, 30 *orb2^RBD1/2*^*, and 9 *orb2^ΔZZ^* brains. Data are pooled from 2 (WT, *orb2^ΔQ^*, and *orb2^ΔZZ^*) or 3 (*orb2^RBD1*^*and *orb2^RBD1/2*^*) independent biological replicates. Significance was determined by Brown-Forsythe and Welch’s ANOVA (H). \**p*<0.05; \*\**p*<0.01; \*\*\**p*<0.001. Scale bars: 40 μm.

The glutamine-rich (poly-Q) domain enables Orb2 to form stable oligomers through prion-like conformational changes that are required for synaptic plasticity, learning, and memory (Si et al., 2003a; Krüttner et al., 2012; Majumdar et al., 2012). Deletion of the poly-Q domain via the *orb2^ΔQ^* allele (Keleman et al., 2007) resulted in brains ∼40% bigger than WT (Fig. 3B,C and G; \*\**p*<0.01 by one-way ANOVA). We used CRISPR-Cas9 editing to mutate both RNA recognition motifs (RRMs), targeting the same amino acids previously identified as critical for RNA-binding activity: Y492, F494, R601, and F604 (Hake et al., 1998; Krüttner et al., 2012). Mutation of RRM1 (Y492A, F494A) or both RRMs (Y492A, F494A, R601P, and F604S; see Methods) was sufficient to reduce brain size (Figure 3D,E and G; *p<0.05 or **p<0.01, respectively, by one-way ANOVA). Within CPEB proteins, the conserved C-terminal cysteine and histidine-rich ZZ domain contributes to protein-protein interactions (Legge et al., 2004; Merkel et al., 2013; Oroz et al., 2020; Low et al., 2025). We analyzed mutants lacking the ZZ domain (AA634-704), *orb2^ΔZZ^*, generated by CRISPR-Cas9. As in *orb2^ΔQ^*, loss of the ZZ domain resulted in brain overgrowth (Fig. 3F and G; \*\**p*<0.01 by one-way ANOVA). The brain growth responses observed in our mutant panel are not due to changes in Orb2 protein expression, as detected by immunoblotting or previously reported (Figure S1F–I) (Xu et al., 2014; Khan et al., 2015). Taken together, these results indicate that the Orb2 RRM1 is necessary to promote brain growth, while the poly-Q and ZZ domains restrict brain overgrowth.

### Orb2-mediated translational activation regulates brain size

Orb2 functions as either a translational repressor or activator depending on its oligomerization status and association with specific cofactors (Fig. 4A) (Si et al., 2003a; Si et al., 2003b; Si et al., 2010; Majumdar et al., 2012; Khan et al., 2015). To determine whether Orb2 regulates brain size through its role in translational control, we depleted its translation cofactors by shRNA (Khan et al., 2015; Stewart et al., 2024). Loss of the repressive cofactor, *CG13928,* did not alter brain volume (Fig. 4B,C; n.s. not significant by two-tailed t-test). However, depletion of the activating cofactor, *CG4612*, increased brain size by ∼30% relative to controls (Fig. 4D,E; \*\*\**p*<0.001 by two-tailed t-test). Our findings that both the Orb2 poly-Q domain and *CG4612* restrict brain growth are consistent with the hypothesis that Orb2 translational activation activity may limit brain size.

**Figure 4.**
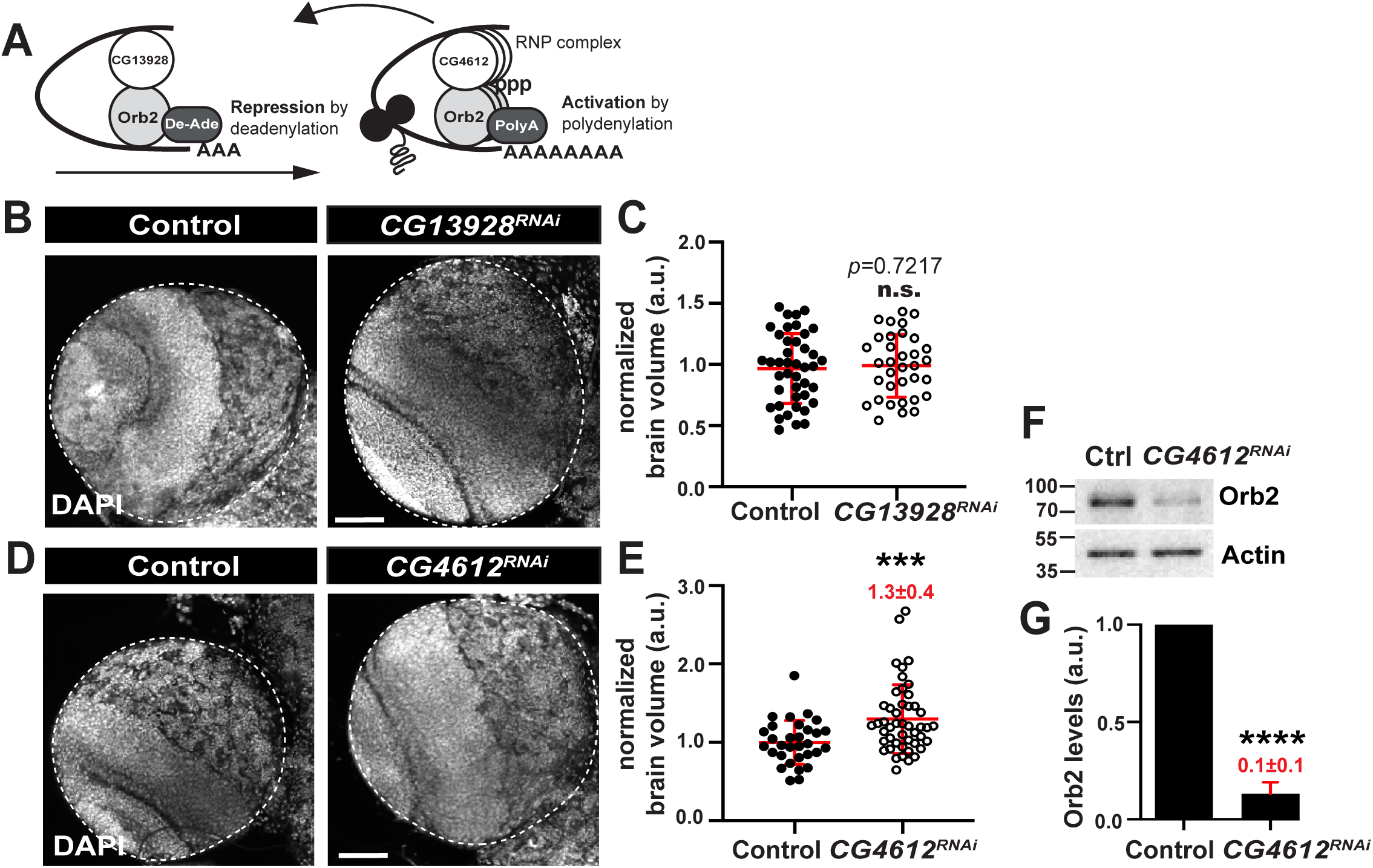
Orb2-mediated translational activation regulates brain size. (A) Schematic of Orb2 repression and activation complexes when bound to cofactors CG13928 or CG4612 (*Khan et. al 2015*). Representative images of age-matched no-GAL4 control and *act-GAL4* driven (B) *CG13928^RNAi^* or (D) *CG4612^RNAi^* third instar larval brains stained for DAPI and associated quantification (C and E), where each dot represents a measurement from a single optic lobe from (C) Control, N= 44 and *CG13828*= 35; (E) Control= 31 and *CG4612*= 47. Data shown are pooled across three replicates. (F) Immunoblot shows levels of Orb2 relative to the actin loading control from control and *CG4612^RNAi^* whole brain extracts, as quantified from triplicated experiments in (G). Significance was determined by Welch’s t-test. n.s., not significant; \*\*\**p*<0.001; \*\*\*\**p*<0.0001. Scale bars: 50 μm. Uncropped blots are available online: https://doi.org/10.6084/m9.figshare.29155505.v1.

Because loss of the translation activation cofactor *CG4612* and *orb2^OE^* both resulted in enlarged brains, we predicted that *CG4612* loss would increase Orb2 protein levels. Unexpectedly, depletion of *CG4612* reduced Orb2 protein levels by ∼90% (Fig. 4F,G; \*\*\*\**p*<0.0001 by two-tailed t-test). These data suggest that CG4612 promotes the translation of Orb2, consistent with Orb2 autoregulation (Majumdar et al., 2012; Stepien et al., 2016). Alternatively, CG4612 may stabilize Orb2. Taken together, we conclude that the Orb2 translational co-activator, CG4612, limits brain growth.

### Orb2 is necessary to maintain the number and size of central brain NSCs

How does Orb2 promote brain growth? To begin to address this question, we tabulated total NSCs in *orb2* versus WT larval brains. Loss of *orb2* or NSC-specific depletion of *orb2* via *wor-GAL4* resulted in ∼60% fewer Mira+ NSCs per lobe relative to WT controls (Fig. 5A,B; *p*<0.0001 by one-way ANOVA). Moreover, as compared to WT, NSC volume was also significantly reduced in *orb2* mutants and following NSC-specific depletion (Fig. 5C; *p*<0.0001 and *p*<0.001, respectively, by one-way ANOVA). These findings are consistent with a requirement of Orb2 in NSC specification and/or maintenance and size.

**Figure 5.**
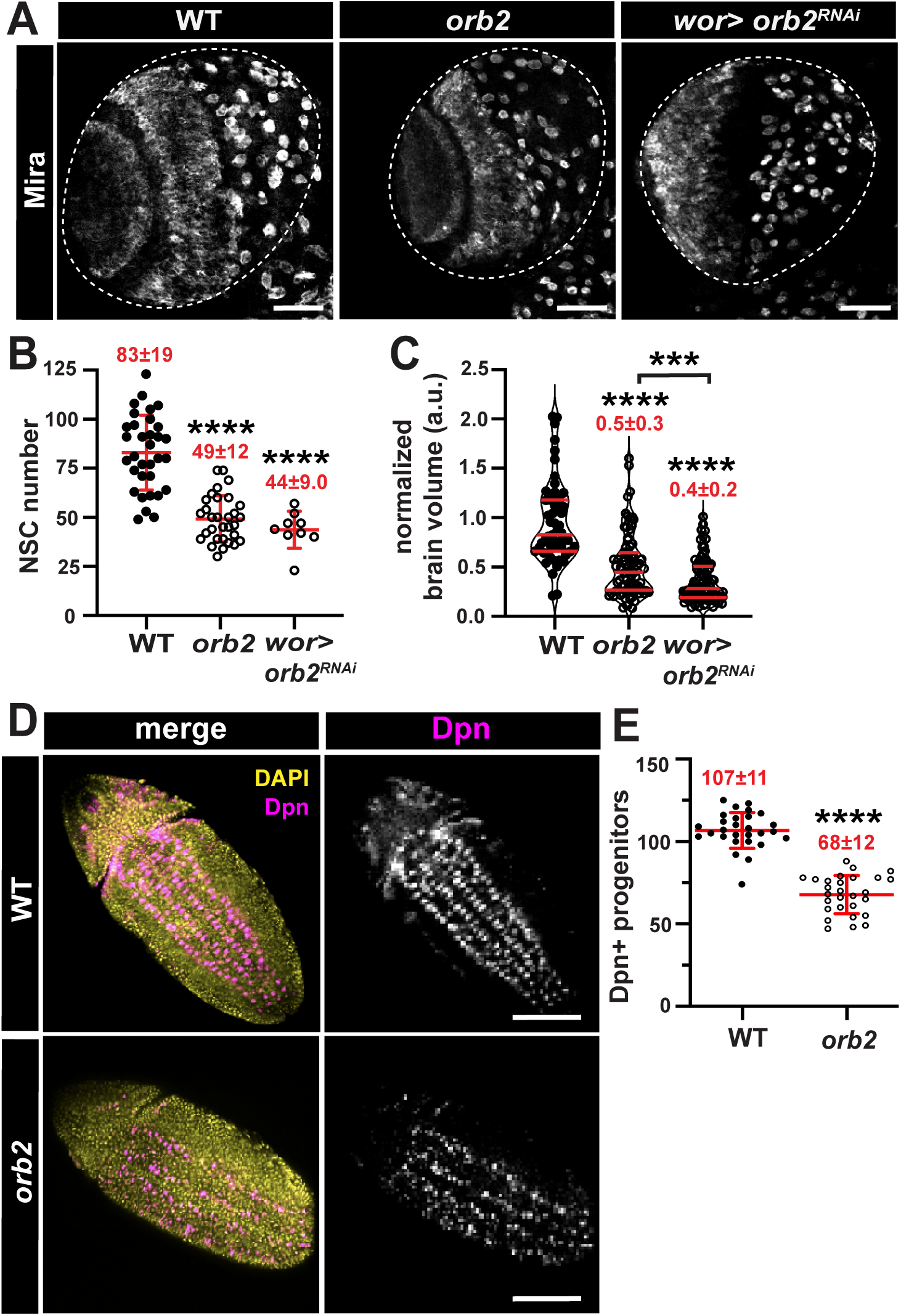
Orb2 enables NSC specification. (A) Representative images show NSCs (Mira, greyscale) within age-matched third instar larval brains of the indicated genotypes. (B) Quantification of NSC number from N= 33 WT, 31 *orb2*, and 9 *wor>orb2^RNAi^*brains. (C) Quantification of NSC volume from WT: N=10 brains and n=55 NSCs, *orb2*: N=10 brains and n=80 NSCs, and *wor>orb2^RNAi^*: N=9 brains and n=90 NSCs. For B and C, each dot represents a measurement from a single optic lobe. (D) Stage 11 embryos stained for Dpn (magenta and greyscale) to label neural progenitors and DAPI (yellow). (E) Quantification of embryonic neural progenitors, where each dot represents a measurement from a single embryo from N=28 WT and *orb2* embryos. Data are pooled from 2 (B and C) or 3 (E) biological replicates. Significance was determined by Brown-Forsythe and Welch’s ANOVA (B and C) or Welch’s t-test (E). \*\*\*\**p*<0.0001. Scale bars: (A) 50 μm and (D) 20 μm.

NSC lineages can be identified based on the expression of defined markers. While all NSCs express the transcription factor Deadpan (Dpn), only Type I NSCs also express Asense (Ase)(Boone & Doe, 2008). The Dpn+ Ase-Type II NSCs produce neurons and glia via intermediate neural progenitors, which divide multiple times to form diverse neural niches (Bello et al., 2008; Boone & Doe, 2008; Izergina et al., 2009). While ∼20% fewer Type II NSCs were noted in *orb2* mutants (*p*<0.01 by one-way ANOVA), no change was detected upon depletion NSC-specific depletion of *orb2* via *wor*-*GAL4*, which expresses in both Type I and Type II NSCs (Ashraf et al., 1999; Weng & Cohen, 2015) (Fig. S2 A,B n.s. by one-way ANOVA). Thus, Type II NSCs may be less sensitive to loss of *orb2* relative to Type I NSCs.

Consistent with a reduction in *orb2* activity, depletion of *CG4612* also decreased total NSCs (Fig. S2 C-E). Intriguingly, the NSCs within *CG4612^RNAi^* brains were enlarged (Figure S2F, G; \*\*\**p*<0.001 by two-tailed t-test). These results suggest that brain size may be uncoupled from NSC number and more closely align with cell size. Taken together, these data imply reduced *orb2* activity may disrupt NSC growth processes, perhaps through the misexpression of cell growth-related proteins.

### Orb2 enables NSC specification

To elucidate mechanisms underlying NSC loss in *orb2* mutants, we first tested the hypothesis that NSCs are eliminated by cell death. We quantified the coincidence of pro-apoptotic cleaved caspase 3 (CC3; (Fan & Bergmann, 2010)) in NSCs marked with the Dpn antibody (Bier et al., 1992; Zhu et al., 2012)). Similar rates of apoptosis were observed in WT and *orb2* NSCs, indicating NSC loss occurs by other mechanisms (Fig. S3 A-C; n.s. by t-test). Analysis of death caspase protein-1 (Dcp-1) also failed to uncover elevated rates of NSC death in *orb2* mutants (Figure S3 D-F). Another mechanism whereby NSCs may be depleted is via premature differentiation (Cabernard & Doe, 2009; Lai & Doe, 2014; Abdel-Salam et al., 2020). Normally, the pro-differentiation marker Prospero (Pros) is confined to the nucleus of the differentiating ganglion mother cell and absent from NSCs (Doe et al., 1991; Vaessin et al., 1991), while retention of Pros in the NSC nucleus promotes premature differentiation (Lai & Doe, 2014). We quantified the coincidence of Pros with Dpn and detected no significant difference relative to WT, indicating premature differentiation does not direct NSC loss in *orb2* mutants (Fig. S3 G-I; n.s. by t-test). Finally, we examined whether *orb2* affects mitotic progression, reasoning impaired NSC self-renewal might contribute to NSC loss. However, the mitotic indices in WT and *orb2* brains were not significantly different (33.6+4.9% per lobe in WT vs. 36.3+10.1% in *orb2* brains; Fig. S3 J-L; n.s. by t-test). Taken together, these data suggest that NSC loss is not due to excess apoptosis, premature differentiation, or impaired proliferation.

Both Type I and Type II NSCs arise from proneural clusters during embryogenesis and serve as the primary source of larval NSCs (Walsh & Doe, 2017; Álvarez & Díaz-Benjumea, 2018). We therefore examined whether *orb2* is required during embryonic neurogenesis for NSC development. Indeed, *orb2* mutant embryos had ∼40% fewer Dpn+ neural progenitors compared to age-matched WT embryos (Fig. 5 D,E; *p*<0.001 by two-tailed t-test). Taken together, these data underscore a role for Orb2 in the specification and/or maintenance of embryonic neural progenitors and suggest that the reduced brain size in *orb2* third instar larvae may result, at least in part, from the reduced number of NSC specified in embryos.

### Orb2 activity is required throughout neurogenesis to regulate brain size

Having identified a requirement for Orb2 in embryos, we next assessed the developmental requirement of *orb2* to promote brain growth. We employed a temperature-sensitive GAL80 mutant (*GAL80^TS^*) to selectively deplete *orb2* in embryos versus larvae. For this strategy, raising progeny at permissive temperatures (18 °C) blocks expression of *orb2^RNAi^*, while growth at the restrictive temperature (29 °C) inactivates *GAL80^TS^*, leading to *orb2^RNAi^* expression and depletion of *orb2* (Fig. 6 A; (McGuire et al., 2003; Zhou et al., 2016)). To specifically deplete *orb2* in early embryos, embryos were reared at the restrictive temperature and then shifted to the permissive temperature. In contrast, the role of *orb2* in larval development was examined by rearing embryos at the permissive temperature until larval hatching, at which point larvae were shifted to the restrictive temperature (Figure 6A, B). Depleting *orb2* during embryogenesis or larval development resulted in significantly reduced larval brain volumes relative to controls (Fig. 6C,D; *p*<0.01 and *p*<0.0001, respectively, by one-way ANOVA). We conclude that Orb2 activity is required throughout neurogenesis to promote brain growth.

**Figure 6.**
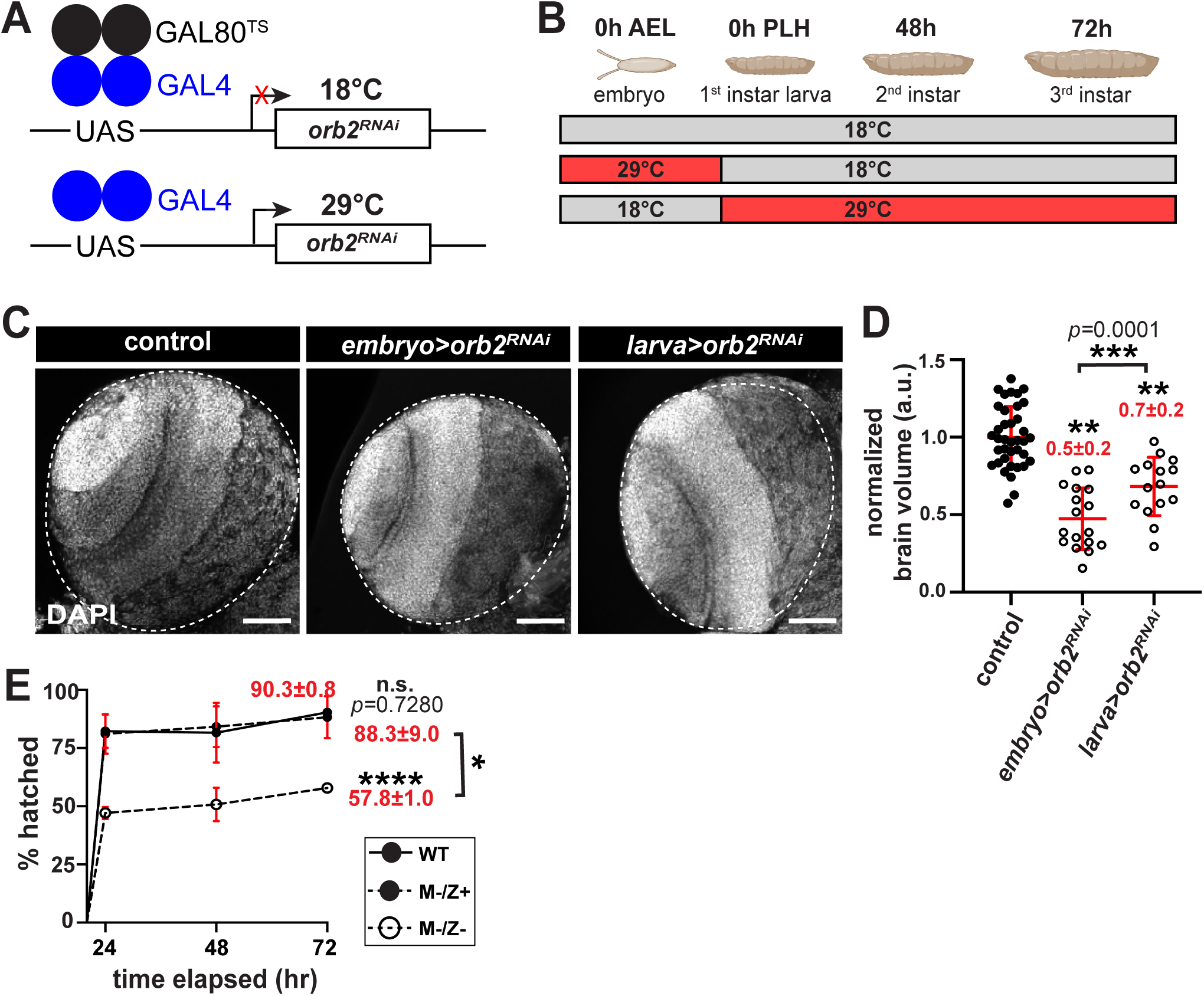
*orb2* regulates brain size throughout neurogenesis. (A) Schematic of the temperature-sensitive depletion of *orb2* by RNAi. (B) The temperature shift schemes used in this study: (top row) controls (*GAL80^TS^, UAS-orb2^RNA^*^i^) were reared at 18°; (middle row) to deplete *orb2* activity in embryos (*embryo*>*orb2^RNA^*^i^), embryos expressing *act>GAL4* and *GAL80^TS^*, *UAS-orb2^RNAi^*were reared at 29°C, then shifted to 18°C 24h PLH; (bottom row) to deplete *orb2* activity in larvae (*larva*>*orb2^RNA^*^i^), embryos expressing the same transgenes were reared at 18° until 24h PLH, then shifted to 29°C. (C) Representative images of age-matched third instar larval brains from controls, embryos raised at the restrictive temperature (*embryo*>*orb2^RNA^*^i^), and larvae raised at the restrictive temperature (*larva*>*orb2^RNA^*^i^), as outlined in (B), and stained for DAPI. (D) Quantification shows normalized third instar brain volume from N=39 control, 17 *embryo*>*orb2^RNA^*^i^, and 15 *larva*>*orb2^RNA^*^i^ brains, where each dot represents a measurement from a single optic lobe. Data shown are pooled from three replicates. (E) Hatch rate analysis of maternal mutant (M-/Z+) versus maternal and zygotic mutant (M-/Z-) *orb2* embryos relative to WT, where each dot represents the average across 4 replicates from N= 800 WT or *orb2* (M-/Z+) and N=820 *orb2* (M-/Z-) embryos. Significance was determined by Brown-Forsythe and Welch’s ANOVA. n.s., not significant; \**p*<0.05; \*\**p*<0.01; \*\*\**p*<0.001; \*\*\*\**p*<0.0001. Scale bars: 50 μm.

To further examine Orb2 requirements during embryogenesis, we conducted a hatch rate analysis. While most WT embryos hatch into first instar larvae within 24-hr, about half of *orb2* null embryos (M^-^/Z^-^) failed to hatch after 72-hr, indicating Orb2 is required for embryonic viability. To distinguish whether *orb2* is maternal-effect lethal or required zygotically, after the maternal-to-zygotic transition, we mated *orb2* virgin females to WT males (Tadros & Lipshitz, 2009; Vastenhouw et al., 2019). The resulting *orb2*/+ (M^−^/Z^+^) embryos hatched at WT levels, indicating that maternal *orb2* is not required for viability. We conclude that *orb2* is necessary zygotically for embryonic survival, and that escaper larvae require *orb2* for proper neurodevelopment (Fig. 6E).

### Orb2 is required for neurogenesis

Our data show that *orb2* loss or depletion results in fewer total NSCs. Because NSCs give rise to the neurons and glia, we also tabulated the number of these cell types within *orb2* mutant larvae relative to controls. Loss of *orb2* resulted in ∼50% more glia, as compared to WT (Fig. S4 A, B; \*\**p*<0.01 by two-tailed t-test). In addition, significantly more *orb2* glia were proliferating relative to controls, which may contribute to their expansion (Pereanu et al., 2005) (Fig. S4A and C; \*\**p*<0.01 by two-tailed t-test). In contrast, neurons were ∼30% reduced in mutants (Fig. S4 D, E; \*\**p*<0.01 by two-tailed t-test). Together, these data suggest that *orb2* normally functions to promote neuronal versus glia cell fate.

### *orb2* is epistatic to the tumor suppressor, *brat*

Brat is a translational repressor and tumor suppressor also implicated in brain size determination. *brat* loss results in massive expansion of Type II NSCs and brain overgrowth (Bello et al., 2006; Betschinger et al., 2006; Lee et al., 2006; Bowman et al., 2008). Considering their opposing phenotypes, we reasoned that Orb2 might instruct brain size by regulating Brat levels. In support of this model, the *brat* 3’UTR contains several cannonical CPE sites and was previously identified as a direct Orb2 RNA target (Fig. 7A and S5 A,B; (Hake & Richter, 1994; Hake et al., 1998; Mastushita-Sakai et al., 2010; Stepien et al., 2016)).

**Figure 7.**
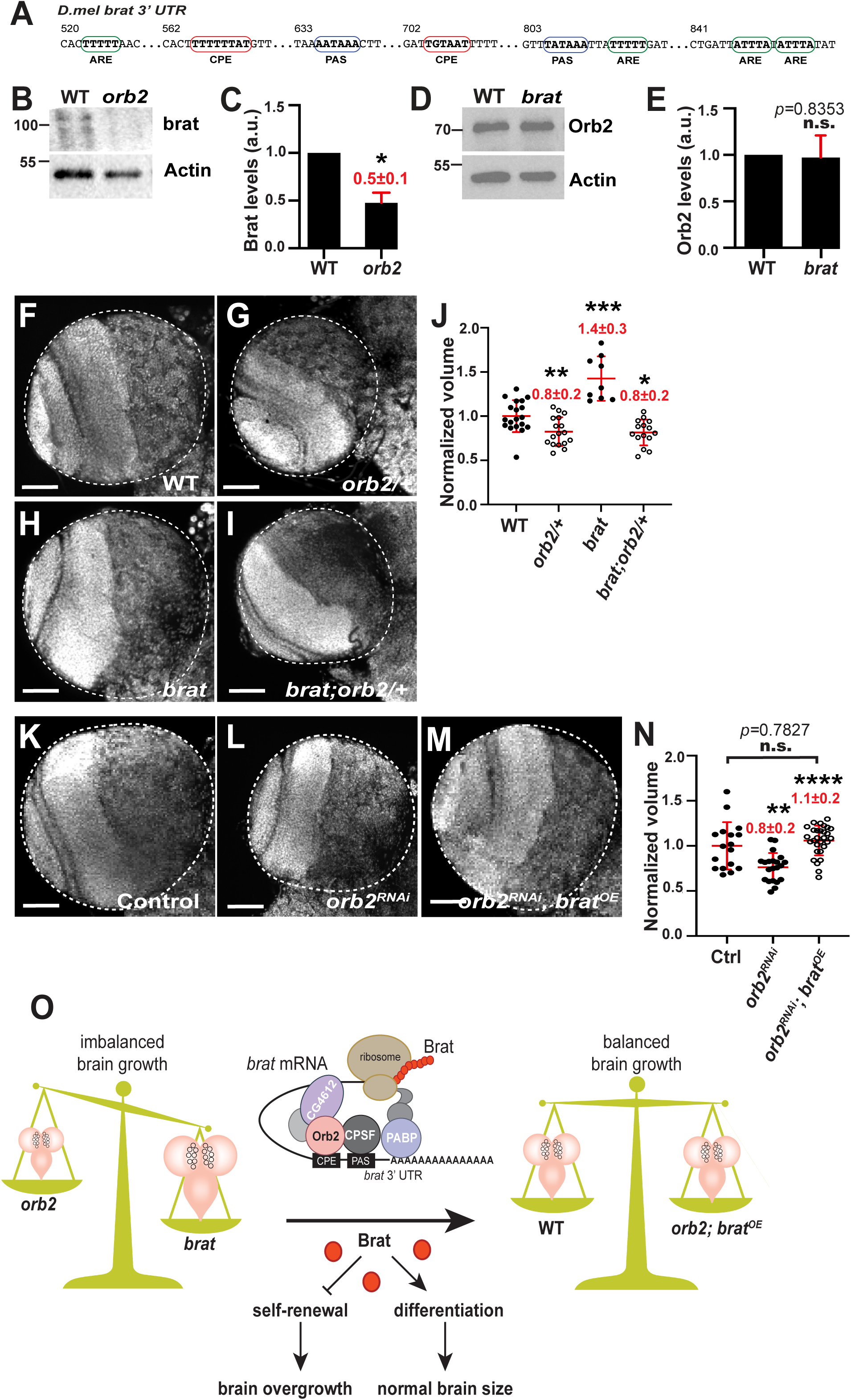
*orb2* is epistatic to *brat*. (A) Schematic of conserved CPE sites in the *brat* 3’UTR. (B) Immunoblot shows Brat protein levels relative to the actin loading control from WT and *orb2* whole brain extracts, as quantified from triplicated experiments in (C). (D) Immunoblot showing Orb2 levels are unchanged in *brat [fs1/k06028]* transheterozygotes relative to WT, as quantified from triplicated experiments in (E). (F–I) Representative images of age-matched third instar larval brains from WT, *orb2/+* hemizygote, *brat [fs1/k06028]*, and *brat;orb2/+* mutants stained for DAPI. (J) Quantification of brain volume normalized to WT, where each dot represents a single measurement from N=19 WT, 17 *orb2/+*, 9 *brat*, and 8 *brat;orb2/+* brains. (K–M) Representative images of age-matched third instar larval brains from *orb2^RNAi^* control, *act>orb2^RNAi^,* and *brat^OE^* rescue (*act>orb2^RNAi^* and *UAS-brat-myc*) stained for DAPI. (N) Quantification of brain volume normalized to control, where each dot represents a single measurement from N= 17 control, 21 *orb2^RNAi^*, and 29 *brat^OE^* brains. Data are pooled from 2 (J) or 3 (N) biological replicates. (O) Proposed model of *orb2*-regulated brain growth through *brat*. Loss of *orb2* results in smaller brains, while loss of *brat* causes brain overgrowth. *orb2* functions upstream of *brat*, likely through regulation of Brat translation. Restoring Brat levels is sufficient to rescue improper brain growth in *orb2* mutants. Significance was determined by Brown-Forsythe and Welch’s ANOVA or Welch’s t-test (E). n.s., not significant; \**p*<0.05; \*\**p*<0.01; \*\*\**p*<0.001; \*\*\*\**p*<0.0001. Scale bars: 50 μm. Uncropped blots are available online: https://doi.org/10.6084/m9.figshare.29155505.v1.

We aligned a region within the *brat* 3′UTR conserved across multiple *brat* isoforms to multiple *Drosophila* species using the conservation insect track on the UCSC genome browser and found that these CPE motifs were conserved across millions of years of evolutionary distance, hinting they may be functionally important (Fig. 7A; (Kent et al., 2002)). To promote translation, CPEB proteins bind CPE sites within 100 nt of polyadenylation signal (PAS) motifs (i.e., the hexanucleotide motif AAUAAA), which recruit the cleavage and polyadenylation specificity factor (CPSF) to cleave pre-mRNA and add the poly(A) tail, aiding mRNA stability, export, and translation (Mendez et al., 2000). We identified PAS motifs (blue boxes) within the *brat* 3′UTR proximal to multipe CPEs (red boxes), consistent with a role for Orb2 in translational activation (Fig. 7A and S5A–C). We further identified multiple AU-rich elements (AREs; green boxes), which modulate mRNA decay or translation, in this same region (Fig. 7A and S5C). This arrangement of CPE, PAS, and ARE motifs suggests precise regulation of *brat* expression to enable normal neurodevelopment (Hake et al., 1998; Mendez & Richter, 2001). These findings also suggest that Brat may be subject to Orb2 translational regulation (Mastushita-Sakai et al., 2010; Santana & Casas-Tintó, 2017). Consistent with this idea, several Orb2 reponse elements (OREs), as recently identified in Orb2 translational targets (Low et al., 2025), are also present in the *brat* 3’UTR, further suggestive of a role in translational regulation.

Orb2 represses the expression of a Brat translation reporter in *Drosophila* S2 cells (Mastushita-Sakai et al., 2010). To uncover the relationship between Orb2 and Brat within larval brains, we first examined their relative distributions. As previously reported, Brat is ubiquitously distributed throughout the CNS. In mitotic NSCs, however, Brat colocalizes with the adapter protein Mira as a crescent at the basal cortex (*arrowheads*, Fig. S5A; (Bello et al., 2006; Betschinger et al., 2006; Lee et al., 2006)). In contrast, Orb2 was undetectible in NSCs or within Mira+ basal crescents (Fig. S5A). Intriguingly, while Orb2 and Brat did partially overlap within differentiated neurons, regions of high Orb2 enrichment contain markedly less Brat, consistent with a potential role for Orb2 in modulating Brat levels (*dashed region*, Fig. S5A).

To assess if Orb2 regulates Brat expression in larval brains, we examined steady-state Brat levels in WT versus *orb2* mutants by western blot. Loss of *orb2* reduced Brat protein levels from larval brain extracts by ∼50% (Fig. 7B,C). By comparison, Orb2 levels are unchanged in *brat* larval brain lysates (Fig. 7D,E). These data are consistent with a role for Orb2 in the translational control of Brat.

We next assayed whether *orb2* and *brat* genetically interact to regulate brain size. Consistent with a strong synthetic lethal interaction, we were unable to recover *brat; orb2* homozygous double-mutant larvae. Nevertheless, we examined whether hemizygosity of *orb2*, previously shown to reduce *orb2* mRNA and protein levels by ∼75% and ∼60%, respectively (Gilmutdinov et al., 2021), was sufficient to modify the *brat* brain overgrowth phenotype. *orb2*/+ hemizygotes showed reduced brain volumes relative to WT, consistent with a dose-dependent effect of *orb2* on brain size (Fig. 7J, F and G; \*\**p*<0.01 by ANOVA). In contrast, loss of *brat* led to the expected increase in brain size (Fig. 7H,J; \*\*\**p*<0.001 by ANOVA). Critically, hemizygosity of *orb2* was sufficient to supress the brain overgrowth of *brat* mutants (*brat*; *orb2*/+; Fig. 7F-J; \**p*<0.05 by ANOVA), indiciating that *orb2* is epistatic to *brat* in the regulation of brain size.

Taken together, our data support a model whereby Orb2 regulates the translation of Brat to modulate brain growth. Such a model predicts that restoring Brat levels in *orb2* mutants would rescue *orb2*-dependent microcephaly. Consistent with this idea, overexpression of a *brat* transgene in an *orb2*-depleted background fully restored brain size (*orb2^RNAi^, brat^OE^*; Fig. 7K-N; \*\*\*\**p*<0.0001). We conclude that Orb2 functions upstream of Brat to balance brain growth (Fig. 7O).

## DISCUSSION

CPEB proteins like Orb2 are important for learning and memory (Si et al., 2003a; Keleman et al., 2007; Krüttner et al., 2012; Majumdar et al., 2012; Khan et al., 2015; Hervás et al., 2016; Li et al., 2016; Oroz et al., 2020; Kozlov et al., 2021; Kozlov et al., 2023). Here, we define a novel and dose-dependent requirement of Orb2 in regulating brain size. The dynamic regulation of *orb2* expression throughout neurogenesis likely ensures proper *orb2* dosage required for neurodevelopment. We find that Orb2 functions in early neural progenitors and *elav*+ neurons to promote brain growth through a mechanism also requiring its RNA-binding activity. We further uncover Orb2 translational activity regulates brain size, as shown by loss of the Orb2 poly-Q domain or the translation co-activator, *CG4612*.

The mechanism by which CPEB proteins regulate target transcripts includes modulating mRNA translation, stability, and localization through CPE elements to influence gene expression. Like other CPEB proteins, the Orb2 RRMs are required for RNA-binding activity (Hake et al., 1998; Krüttner et al., 2012; Stepien et al., 2016). However, we unexpectedly uncovered opposing phenotypes in response to deleting different Orb2 functional domains. While the RRMs support brain growth, consistent with the *orb2* null phenotype, the poly-Q and ZZ domains apparently restrict overgrowth. Given that Orb2 autoregulates its own translation via CPE motifs in its 3’ UTR, the various Orb2 domains may play conformationally significant and conserved roles in forming complexes with cofactors that influence expression of neurogenic genes or pro-growth factors (Tan et al., 2001).

Given the varying phenotypes resulting from our structure-function analysis, we speculated that Orb2 may promote the translation of a repressor of brain growth. Such a factor would act in opposition to Orb2, which normally promotes brain growth. This model would establish a feedback loop to normalize brain growth, consistent with a role for Orb2 in the homeostatic regulation of brain size.

Our data suggest that Orb2 regulates brain size through translational regulation of *brat* mRNA (Figure 7O). This model is supported by prior work demonstrating that Orb2 binds *brat* mRNA directly at defined CPE sites within the *brat* 3’UTR (Stepien et al., 2016). We further identify hexanucleotide CPSF binding sites proximal to conserved CPE sites necessary for CPEB-dependent translational activation (Mendez et al., 2000). We also show Orb2 regulates Brat protein levels. Finally, we demonstrate that *orb2* is epistatic to *brat* and that restoring Brat levels by expression of a *UAS-brat* transgene is sufficient to rescue *orb2*-dependent microcephaly.

In the early embryo, Brat represses *hunchback* (hb) translation, critical for axial patterning (Sonoda & Wharton, 2001). In neurogenic embryos, Brat segregates asymmetrically with Mira in dividing NSCs to promote neuronal differentiation (Betschinger et al., 2006). In larvae, Brat limits NSC proliferation by regulating various transcription factors, including dMyc, and works alongside Pros to repress NSC renewal and promote differentiation (Betschinger et al., 2006).

While our study suggests Orb2 regulates Brat levels to ensure proper NSC specification, growth, and neurogenesis, other factors must cooperate with Orb2 to tune Brat expression. *orb2* loss reduced, but did not eliminate, Brat. Likewise, we did not detect changes in NSC proliferation in *orb2* mutants. Further study is necessary to identify additional regulators of *brat* activity and how such factors coordinate with Orb2.

Interestingly, temperature-shift experiments uncovered requirements for *orb2* function during both the embryonic and larval neurogenic stages. Indeed, our data suggest that Orb2 may be required for NSC specification in embryos. This early requirement for *orb2* likely establishes the NSC pool required at later developmental stages. During normal development, late embryonic NSCs enter quiescence. In early larval stages, however, NSCs reactivate, an early step of which involves the doubling in NSC size (4–5 µm to 10–15 µm; (Truman & Bate, 1988; Prokop & Technau, 1991; Ding et al., 2016)). Consequently, the dividing NSCs normally produce daughter cells with marked cell size asymmetries: a larger, self-renewing stem cell and a smaller daughter cell fated for differentiation. In addition to being reduced in number, *orb2* NSCs are also smaller. These phenotypes contrast with *brat* mutant NSCs, which fail to inhibit self-renewal in the daughter cell. Instead, both cells grow and proliferate, leading to brain tumors (Betschinger et al., 2006; Lee et al., 2006). Taken together, these findings suggest that Brat and Orb2 likely have distinct functions in NSCs versus mature neurons.

One way in which Brat regulates cell growth is through coordination of ribosome biogenesis and translational control (Gui et al., 2023). One possibility raised by our study is that *orb2* may be required for NSC growth during NSC reactivation. NSC reactivation requires feeding and nutrient sensing (Britton & Edgar, 1998). Because we used an age-matching paradigm dependent upon larval intake of blue-colored food throughout this study (see Methods and (Hailstock et al., 2023)), any *orb2*-dependent response to NSC reactivation must occur post-feeding. We speculate that Orb2 functions in either the transmission of the cell growth signals themselves or in the physical enlargement of the NSCs. These processes may involve the recently defined role for Orb2 in enabling the translation of rare-codon-enriched mRNAs (Stewart et al., 2024). Further investigation is required to elucidate precisely how Orb2 influences NSC specification and growth.

## MATERIALS AND METHODS

### Fly stocks

The following strains and transgenic lines were used: *y^1^w^1118^*(Bloomington *Drosophila* Stock Center (BDSC) #1495) was used as the WT control unless otherwise noted. *orb2* brains were isolated from the homozygous null allele, *orb2^Δ36^* (P. Schedl, Princeton University (Xu et al., 2012). *orb2* null embryos were sorted against a GFP balancer (BDSC #58479). Tissue-specific depletion of *orb2* was by *orb2^RNAi^* (P{TRiP.JF02376}attP2; BDSC #27050). *UAS-mCherry_NLS_*} (BDSC #38425) was used to visualize GAL4 expression pattern, and *orb2[+GFP]* (BDSC #94349; gift from K. Keleman, (Krüttner et al., 2012)) is a GFP knock-in expressing *Orb2-GFP* at the native *orb2* locus and with endogenous *orb2* regulatory elements was used to detect Orb2 localization. A *UAS-orb2B-GFP* transgene was used in overexpression experiments (gift K. Keleman, (Krüttner et al., 2012)). UAS transgenes were driven by *wor-GAL4* (P{wor.GAL4.A}2; BDSC #56555), *insc-GAL4* (P{GawB}insc^Mz1407^; BDSC #8751), *elav-GAL4* (P{w[+]elav.GAL4}), *elav^c155^-GAL4* (P{GawB}elav^C155^; BDSC #458), *repo-GAL4* (P{GAL4}repo; BDSC #7415), *c855a-GAL4* (P{GawB}c855a; BDSC #6990), *brp-GAL4* (gift from A.-S. Chiang and K.-L. Feng, National Tsing Hua University; (Chen et al., 2023)), or *act-GAL4* (P{w[+mC]=Act5C-GAL4}; BDSC #4414). Temperature-sensitive GAL80 (P{GAL80^ts^}10; BDSC #7108) was recombined with *orb2^RNAi^. orb2^ΔQ^* was used to assess functional requirement of the poly-Q domain (gift from P. Schedl, Princeton University; (Keleman et al., 2007). *CG4162^RNAi^* (P{TRiP.HMJ22843}attP40; BDSC #60473) and *CG13928^RNAi^* (P{TRiP.HM05031}attP2; BDSC #28545) were used for cofactor depletion. *brat^fs1^* and *brat^k06028^* (gifts from C.Y. Lee, University of Michigan; (Arama et al., 2000)) were used to generate *brat^fs1^*/*brat^k06028^*trans-heterozygote animals. *brat^k06028^* is a null allele (Sonoda & Wharton, 2001). *brat^fs1^* is a loss-of-function allele defined by a G774D mutation in the NHL domain (Sonoda & Wharton, 2001) and was recombined onto the *orb2* chromosome to generate *brat^fs1^, orb2^Δ36^* recombinant animals. *UAS-brat^FL^:myc* (gift from C.Y. Lee, University of Michigan; (Komori et al., 2014) was recombined with *orb2^RNAi^* to overexpress *brat*. All strains were maintained on Bloomington formula cornmeal-agar media (Lab-Express, Inc.; Ann Arbor, MI) at 18, 25, or 29°C in light and temperature-controlled incubators.

### Generation of *orb2* mutant alleles

*orb2^RBD1*^* and *orb2^RBD1/2*^* mutants were generated by CRISPR using the Scarless approach (Bier et al., 2018) using the method of (Port & Bullock, 2016). Multicistronic target amplicons were generated by PCR and inserted into BbsI-digested pCFD5 (Addgene #73914) by Gibson assembly using the following 5’–3’ forward (F) and reverse (R) primers with gRNA underlined:

1. F: GCGGCCCGGGTTCGATTCCCGGCCGATGCA<u>CAGCAGGAAGGCATATCCCT</u>GTT TTAGAGCTAGAAATAGCAAG and
2. R: ATTTTAACTTGCTTTTCTAGCTCTAAAAC<u>CGTAGTCAGCCAATTTCCTC</u>TGCACC AGCCGGGAATCGAACCC

For homologous recombination, gene blocks were synthesized (Twist Bioscience) then sequentially ligated into pHD-DsRed-Scarless (*Drosophila* Genomics Resource Center (DGRC), #1364). DNA plasmids were injected by BestGene (Chino Hills, CA) into attP40 *nos-Cas9* (BDSC#78781 or TH00788.N). The PAM sequences at both target cut sites were mutated with silent mutations to prevent recutting. Two unique CRISPR lines were recovered and sequence validated using primers:

1. orb2RRMF: GCTGGGTGGAAATGGTGG and
2. orb2RRMR: GCGTGTGTTGTCTCTAATGGT

Alanine substitutions were designed for four amino acids (two in each RRM), following the design of (Krüttner et al., 2012). One isolate had mutations at all expected sites despite alternate amino acid substitutions in RRM2: Y492A, F494A, R601P, and F604S. One mutant had mutations only in RRM1: Y492A and F494A. Mutant lines were backcrossed to *alphatub-piggyBacK10* (BDSC #32070) to excise the DsRed marker, then maintained over the TM6C, Sb,Tb balancer chromosome.

CRISPR mutant *orb2^ΔZZ^* was generated according to the method of (Gratz et al., 2014) as described in (Low et al., 2025). The *orb2^ΔZZ^*mutation deletes AA634-704 spanning the ZZ domain of Orb2B, and the CRISPR mutants were screened and confirmed by DNA sequencing.

### Hatch rate analysis

Embryos from *y^1^w^1118^* control, *orb2^Δ36^*, and heterozygotes were collected on yeasted grape juice agar plates and ∼200 embryos were transferred at 0-2h AEL to fresh plates and aged for 72 h at 25°C. Unhatched embryos were counted from each plate and compared to controls. *orb2^Δ36^* virgins and males were mated to generate M-Z-embryos, while *orb2* heterozygotes (M-Z+) were derived by crossing null virgin females to control males, to assess maternal versus zygotic contributions. Data presented are mean ± SD from three biological replicates.

### Larval age matching

Larval samples were synchronously staged as described by (Hailstock et al., 2023). Briefly, *Drosophila* adults were allowed to seed grape juice agar plates supplemented with yeast paste for 48 h at 25°C. Plates were changed every 24 h, after which flies were allowed 30 minutes to clear previously fertilized eggs. Embryos were collected for 4 h on a fresh plate and incubated at 25°C until first instar larvae hatched. First instar larvae were transferred into vials containing *Drosophila* culturing medium supplemented with yeast paste dyed with bromophenol blue, then incubated at 25°C until crawling third instar larvae emerged. Age-matched larvae that both displayed foraging behavior and partial gut clearance of the blue yeast paste were then selected for dissection (Truman & Bate, 1988).

### Temperature-shift assay

For temperature-shift experiments, we used *GAL80^TS^* to modulate expression of *orb2^RNAi^*. Animals were reared at 18 °C to inhibit *orb2^RNAi^* expression or at 29 °C allow *orb2^RNAi^*expression. *act-GAL4* virgins were mated to *GAL80^TS^, orb2^RNAi^* recombinant animals in cages for 48 h at 18°C. At 24-48h AEL, *orb2* expression was inhibited in first instar larvae by shifting animals to 29°C. To inhibit *orb2* activity in embryos, the same parental genotypes were mated at 29°C for 48 h prior to harvesting 4-7 h embryos. Embryos were maintained at 29°C for 24-48h, and first instar larvae were shifted to 18°C to allow *orb2^RNAi^* expression. Age-matched third instar larvae were selected 48-72 h after temperature-shift as described in (Hailstock et al., 2023). Experimental groups were compared to *GAL80^TS^, orb2^RNAi^* controls lacking GAL4 raised at 18°C.

### Immunofluorescence

Age-matched crawling third instar larval brains were used for dissections and prepared for immunofluorescence as previously described (Lerit et al., 2014). Brains were dissected in Schneider’s *Drosophila* Medium (ThermoFisher Scientific, #21720024), fixed in 9% paraformaldehyde for 15 min, blocked in PBT buffer [Phosphate Buffered Saline (PBS) supplemented with 1% bovine serum albumin (BSA) and 0.1% Tween-20] for one hour at room temperature prior to overnight incubation in primary antibodies in PBT with nutation at 4°C. Samples were further blocked with modified PBT (2% BSA, 0.1% Tween-20, and 4% normal goat serum (NGS) in PBS) for one hour before incubation for two hours at room temperature with secondary antibody and DAPI.

Embryos were fixed in a 1:4 solution of 4% paraformaldehyde:heptane for 20 min and devitellinized in methanol (Rothwell & Sullivan, 2007). Fixed embryos were rehydrated, blocked in PBTx buffer (PBS supplemented with 0.3% Triton X-100 and 1% BSA), and incubated overnight at 4°C with primary antibodies diluted in PBTx supplemented with 2% NGS. After washing, embryos were further blocked in PBTx supplemented with 2% NGS and incubated for 2 h at room temperature with secondary antibodies and DAPI (10 ng/ml, ThermoFisher Scientific, #D1306). All samples were mounted in Aqua-Poly/Mount (Polysciences Inc., #18606-20) prior to imaging.

The following primary antibodies were used: guinea pig anti-Orb2 (1:1000, gift from K. Si, Stowers institute for Medical Research; (Mastushita-Sakai et al., 2010)), mouse anti-elav 9F8A9 (1:500, DSHB), mouse anti-Orb2 2D11 and 4GS (1:25 each, DSHB), mouse anti-repo 8D12 (1:2000, DSHB), mouse anti-Prospero MR1A (1:500, DSHB), rabbit anti-DCP-1 (1:500, Cell Signaling Technology #9578), rabbit anti-Dpn (1:500, gift from Y.N. Jan, University of California San Francisco; (Bier et al., 1992)), rabbit anti-pH3 (1:1000, Millipore #05570), and rat anti-Mira (1:500, Abcam, #ab197788). For embryos, guinea pig anti-Dpn (1:500, gift from J. Skeath, Washington University of St. Louis; (Skeath et al., 2017)) was used. Secondary antibodies were Alexa Fluor 488, 568, or 647 (1:500, Molecular Probes) and DAPI (10 ng/ml, Invitrogen).

### Microscopy

Images were acquired on a Nikon Ti-E system fitted with a Yokagawa CSU-X1 (Yokogawa Corp. of America) spinning disk head, Hamamatsu Orca Flash 4.0 v2 digital complementary metal oxide-semiconductor (CMOS) camera (Hamamatsu Corp.), Perfect Focus system (Nikon), and Nikon LU-N4 solid state laser launch (15 mW; 405, 488, 561, and 647 nm) The following Nikon objectives were used: 100x 1.49-NA Apo Total Internal Reflection Fluorescence oil immersion, 40x 1.3-NA Plan Fluor oil immersion, and 20x 0.75-NA Plan Apo. Images were acquired at 25°C through Nikon Elements AR software on a 64-bit HP Z440 workstation (Hewlett-Packard).

### Live Imaging

Age-matched larval brains were dissected as described in (Lerit et al., 2014) and prepared for live imaging as described in (Hannaford et al., 2019). Briefly, explanted brains were transferred to a drop of fibrinogen-supplemented media (10mg/mL fibrinogen (Sigma Aldrich #341573) added to Schneider’s *Drosophila* Medium (ThermoFisher Scientific #21720024)) in a glass bottom imaging dish (MatTek #P35G-1.5-14-C). The brain was oriented with the dorso-anterior central brain contacting the glass, and the clot was polymerized by the addition of thrombin at 100µg/mL (Sigma Aldrich #T7513). After polymerization, 500 μL of medium was added on top of the clot for the duration of imaging.

### Image Analysis

Images were resolved using Fiji (National Institutes of Health; (Schindelin et al., 2012)), to separate or merge channels, crop regions of interest, generate maximum-intensity projections, and adjust brightness and contrast. Adobe Photoshop and Adobe Illustrator software (Adobe, Inc.) were used to assemble images into panels.

### Larval brain volumetric analysis

Volume measurements were obtained in Imaris 9.9, 10.1.0 and 10.1.1 software (Oxford Instruments) using the 3D surface tool to define the optic lobe region of interest on DAPI-visualized brains and statistics function to calculate surface volume. Brain volume is represented by measurement from one lobe per brain and normalized to the control mean (Link et al., 2019). NSC and neuroepithelium volume measurements were obtained using the 3D surface tool to detect objects with machine learning function to subtract background. Cell volume was calculated using the statistics function. The experimenter was blinded to genotype prior to analysis.

### Cell number and volume

For embryos, 4-7 h WT and *orb2* embryos were collected and stained with anti-Dpn antibodies. Maximum projected images of Dpn+ cells were counted in Fiji using the Cell Counter (O’Brien et al., 2016) plugin. The experimenter was blinded to genotype prior to analysis.

For larvae, NSCs were identified as large, Mira+ cells located in the central brain. NSC counts are reported from a single optic lobe per sample. Mitotic index was calculated as the number of Mira+, pH3+ NSCs per total Mira+ NSCs. Mitotic events were identified by the presence of pH3 signal. Single-channel .tif raw images for mitotic event, glia, and neurons were segmented in three dimensions using Python scripts adapted from the Allen Institute for Cell Science Cell Segmenter (Chen et al., 2020) as described in (Ryder & Lerit, 2020). To score cell death, events were identified by the presence of Dcp-1 in Mira+ cells using the Cell Counter (O’Brien et al., 2016) plugin in Fiji. The experimenter was blinded to genotype prior to analysis.

To quantify neurons per brain lobe, age matched WT and *orb2* third instar larvae expressing *elav^C155^-GAL4* and *UAS-mCherry_NLS_* were live imaged. Images were processed with Imaris 9.9 using the spot tool to detect objects and the number of objects were counted with the statistics function. The experimenter was blinded to genotype prior to analysis.

### Single cell RNA-seq analysis

Single cell RNA sequencing data was downloaded from GEO Dataset GSE135810. Count matrices were imported into R, filtered, and quality controlled as described in the Github repository companion to the original publication (Corrales et al., 2022) using SeuratV5 (Hao et al., 2024). Cells were grouped into 67 clusters and then assigned a cell type by comparing the positively differentially expressed genes to known biomarkers as defined by Corrales et al. Supplementary Spreadsheet 19. Visualization was completed in R using Seurat visualization functions.

### Immunoblotting

Larval brain extracts were prepared from 10-20 crawling third instar larva dissected in Schneider’s *Drosophila* Medium, removed of imaginal discs, transferred to fresh media, then rinsed once in cold PBST (PBS with 0.1% Tween-20). Samples were homogenized on ice in 30 μL of fresh PBST using a cordless motor and plastic disposable homogenizer, supplemented with 5X SDS loading buffer to 2X final concentration, boiled for 10 minutes at 95 °C, then stored at −20 °C or immediately resolved on a commercial 10% polyacrylamide gel (Bio-Rad, #4568023). Protein concentration was measure using the Pierce BCA Protein assay kit (Thermo Scientific, #23227) as per manufacturer’s instructions and measured on Synergy H1 Multi-Mode plate reader. Proteins were either transferred to a 0.2 μm nitrocellulose membrane (Amersham Protran, #10600001) or PVDF membrane (Bio-Rad, #1620177) by wet transfer in a buffer containing 25 mM Tris-HCl pH 7.6, 192 mM glycine, 10% methanol, and 0.02% SDS at 4 °C or semi-dry transfer by Trans-Blot Turbo Transfer System (Bio-Rad). Membranes were blocked in 3% BSA or 5% dry milk in TBST (Tris-buffered saline, 0.05% Tween-20), washed with TBST, and incubated overnight at 4 °C with primary antibodies. After washing with TBST, membranes were incubated for 2 h in secondary antibodies diluted 1:2500 in TBST. Bands were visualized with Clarity ECL substrate (Bio-Rad, #1705061) on a ChemiDoc imaging system (Bio-Rad). Densitometry was measured in ImageJ and protein levels were normalized to the loading control as described in (Stael et al., 2022).

The following primary antibodies were used: mouse anti-Actin JLA20 (1:1000, DHSB), mouse anti-Brat 3A9 (1:1000, gift from W.Shi, Institute of Genetics and Developmental Biology, China; (Shi et al., 2013)), and mouse anti-Orb2 2D11 and 4G8 (1:25 dilution each; DSHB). Secondary antibodies: goat anti-mouse HRP (1:2500 or 5000, ThermoFisher #31430) and goat anti-rabbit HRP (1:5000, ThermoFisher # 656120). Densitometry was measured in Adobe Photoshop and protein levels are normalized to the loading control.

### Statistical methods

Data were plotted and statistical analysis was performed using GraphPad Prism software. Data were first tested for any outliers using a ROUT test with a Q= 1% and normality using the D’Agostino and Pearson normality test. This was followed by Student’s two-tailed t test, ANOVA, Fisher’s exact test, or the appropriate nonparametric tests. Data were plotted with mean ± SD displayed. Data shown are representative results from at least two independent replicate experiments as indicated in the figure legends.

## Data availability

All code for object detection and measurements (Fig. S4 A-C, G-L) are available on Github at https://github.com/pearlryder/rna-at-centrosomes (Ryder & Lerit, 2020; Lucas et al., 2021). Full-size, replicated immunoblots are available to view on FigShare: https://doi.org/10.6084/m9.figshare.29155505.v1.

## Acknowledgments

We thank Drs. Ann-Shyn Chiang, Yuh Nung Jan, Cheng-Yu Lee, Tzumin Lee, Wenwen Shi, James Skeath, and Noah Sokol for gifts of reagents and stocks. We are grateful to Dr. Victor Faundez, Dr. Kenneth Moberg, and members of our laboratory for constructive feedback. We thank Drs. Sergei Bombin and Adam Gracz for technical assistance with scRNA-seq analysis. Stocks obtained from the Bloomington *Drosophila* Stock Center (NIH P40OD018537) were used in this study. Resources from the *Drosophila* Genomics Resource Center, supported by NIH grant 2P40OD010949, were used in this study. Antibodies from the Developmental Studies Hybridoma Bank, created by the National Institute of Child Health and Human Development of the National Institutes of Health and maintained at the University of Iowa Department of Biology, Iowa City, IA 52242 were used in this study. Research reported in this publication was supported in part by the Emory University Integrated Cellular Imaging Core of the Winship Cancer Institute of Emory University and NIH/NCI under award number, 2P30CA138292-04, (RRID:SCR_023534).

This work was supported by NIH grants T32GM008367-30 (TH and JB), F31NS134380 (JB), T32GM008490 (BVR), R01GM138544 (DAL), R35GM158404 (DAL), and NIH/NIGMS 1R35GM155195-01 (TAS), and Canadian Institutes of Health Research grants PJT-159702 and PJT-190124 (HDL). TCHL was supported in part by a Canadian Graduate Scholarship (CGS-M) and a University of Toronto Open Fellowship. TH is supported by an HHMI Gilliam Fellowship GT16518. DAL is also supported by a Research Scholar Grant (RSG-22-874157-01-CCB) from the American Cancer Society. The content is solely the responsibility of the authors and does not necessarily represent the official views of the National Institutes of Health or other funders.

## Data Availability Statement

Uncropped immunoblots are available online: https://doi.org/10.6084/m9.figshare.29155505.v1.

## Online supplemental material

Five supplemental figures accompany this manuscript.

## SUPPLEMENTAL FIGURE LEGENDS

**Supplemental Figure 1.**
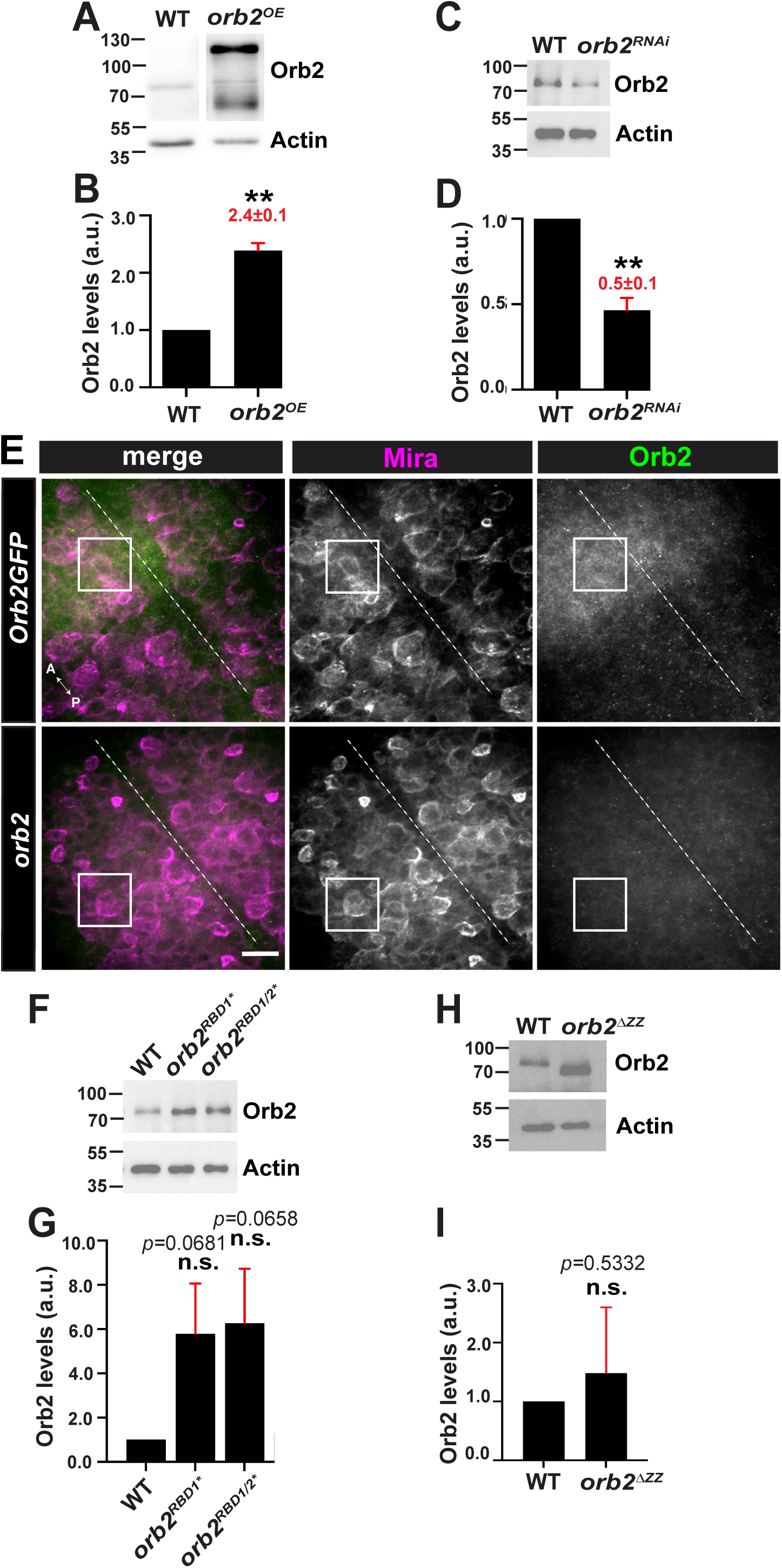
O**r**b2 **expression and localization.** (A) Orb2 expression was quantified by immunoblot from WT and *UAS-orb2-GFP* (*orb2^OE^*) whole brain extracts using anti-Orb2 antibodies relative to the actin loading control, and quantified by densitometry analysis from biological triplicate in (B). (C) *orb2*^RNAi^ efficacy was validated by western blot and quantified in biological triplicate in (D). (E) Maximum intensity projections of a stage 11 *Orb2-GFP* embryo from 4–7 hr collections stained with anti-GFP (green) and anti-Mira antibodies (magenta) compared to *orb2* null. Boxed areas show dorsoanterior neural progenitors (Mira+) within age-matched embryos. Orb2-GFP signals were coincident with Mira in 15/18 embryos. Dashed line indicates the ventral midline. Anterior (A) is to the left, and posterior (P) to the right. (F) Immunoblot of Orb2 protein levels in the indicated genotypes, as quantified from biological triplicate in (G). Significance was determined by Welch’s t-test (**B,D,G,I**). \**p*<0.05; \*\**p*<0.01. Scale bar: 20 µm. Uncropped blots are available online: https://doi.org/10.6084/m9.figshare.29155505.v1.

**Supplemental Figure 2.**
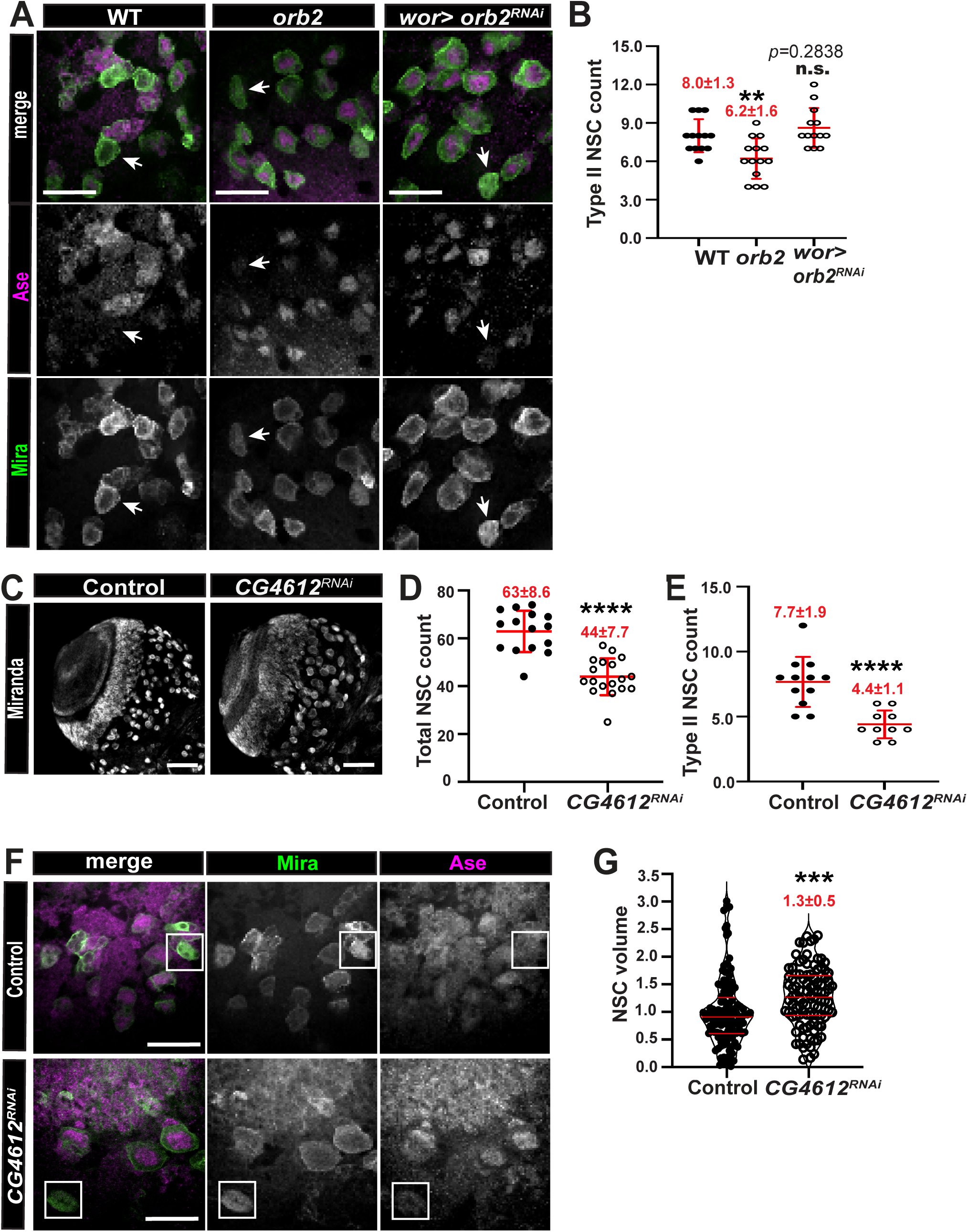
o*r*b2 maintains the NSCs. (A) Maximum intensity projected images of Type II NSCs (arrows) from the indicated genotypes labeled with Dpn (magenta) and Ase (green) antibodies. (B) Quantification of the number of Type II NSCs from N= 13 WT, 14 *orb2*, and 13 *wor>RNAi* third larval instar brains. (C) Images show total NSCs, marked with anti-Mira antibodies, in controls and following depletion of *CG4612.* (D) The total number of Mira+ NSCs was counted from N= 15 control and 18 *CG4612^RNAi^* brains. (E) The number of Type II (Mira+, Dpn-) NSCs from N= 9 control and 6 *CG4612^RNAi^* brains. Data shown in B, D, and E are pooled across two biological replicates, and each dot represents a measurement from a single brain. Mean ± S.D. is shown. (F) Maximum intensity projected control and *CG4612^RNAi^* third instar larval brains stained for Mira (green) to label all NSCs and Ase (magenta) to distinguish Type I vs. Type II NSCs. (G) NSC volume was measured across all Mira+ cells from N=7 control and N= 6 *CG4612^RNAi^* brains and is enlarged following depletion of *CG4612*. Significance was determined by Welch’s t-test. Mean ± S.D. is displayed. n.s., not significant; \*\**p*<0.01; \*\*\**p*<0.001; \*\*\*\**p*<0.0001. Scale bars: (A and F) 20 µm.

**Supplemental Figure 3.**
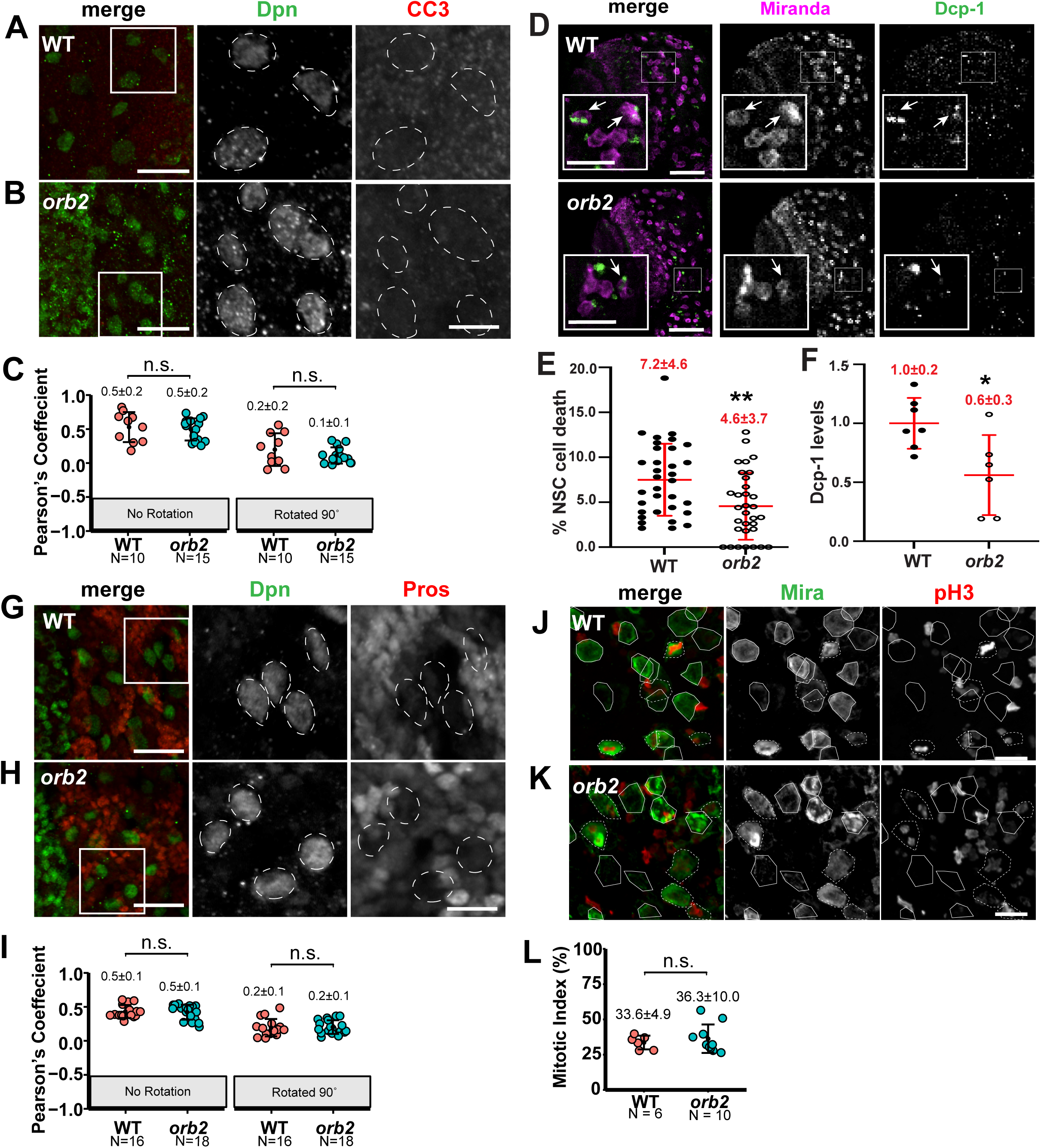
N**S**C **viability and proliferation are not impaired in *orb2* mutants.** Maximum intensity projections of (**A**) WT and (**B**) *orb2* brains stained with Dpn (green; NSC nuclei) and CC3 (red; pro-apoptotic marker). Insets (boxes) are enlarged to the right to highlight NSCs (dashed ovals). (**C**) Quantification of the Pearson’s correlation coefficient between Dpn and CC3 shows non-significant co-incidence from N=10 WT and 15 *orb2* mutant brains. The CC3 channel was rotated clockwise (90°) to test for specificity of overlapping signals. Data shown are representative from two independent biological replicates. (D) Representative third instar larval brains with zoomed insets (boxes) to assay colocalization of Mira (NSCs, magenta) and Dcp-1 (green) cell death events (arrows). (E) Quantification of Mira+ and Dcp-1+ NSC cell death events within the central brain region from N=33 WT and 22 *orb2* brains. Data shown are pooled from two independent biological replicates. Maximum intensity projections of (G) WT and (**H**) *orb2* brains stained with Dpn (green) and Pros (red; differentiated nuclei) to assay premature differentiation. (**I**) Quantification of the Pearson’s correlation coefficient between Dpn and Pros show no significant coincidence from N=16 WT and 18 *orb2* brains. The Pros channel was rotated clockwise (90°) to test for specificity of overlapping signals. Data shown are representative from two independent biological replicates. In (C) and (I), each data point is from one ROI per optic lobe. Images show (J) WT and (**K**) *orb2* NSCs (Mira, green; circles) and the mitotic marker pH3 (red). (**L**) Quantification of mitotic index from N= 6 WT and 10 *orb2* brains. Data shown are representative from two independent biological replicates. Significance was determined by Student’s t-test. n.s., not significant; \**p*<0.05; \*\**p*<0.01. Scale bars: (A,B; G,H) 20 μm and 8 μm (insets); (**D**) 50 μm and 20 μm (insets); and (J and K) 10 μm.

**Supplemental Figure 4.**
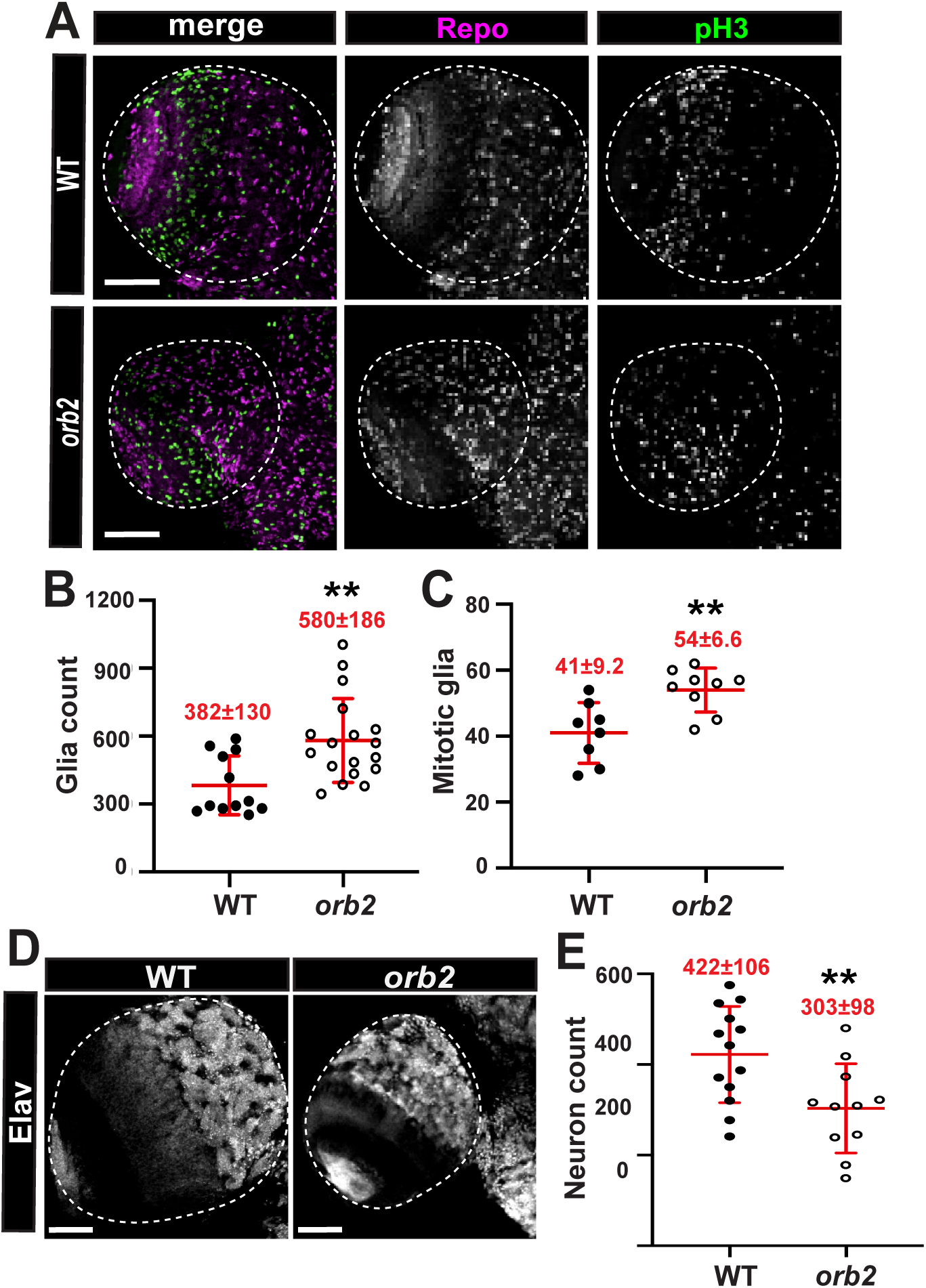
Orb2 balances neural versus glial differentiation in larval brains. (A) Images show larval brains stained for Repo (magenta) to label glia and the mitotic marker phospho-Ser10 Histone H3 (pH3; green) in WT vs *orb2* mutants. (B) More glia cells are detected in *orb2* (N=18 brains) central brain regions, as compared to WT (N=13 brains). (D) images show larval brains stained for Elav (grey) to mark neurons. (E) Fewer neurons are detected from N=11 *orb2* brains relative to N=13 WT brains. Data shown are pooled across two biological replicates. Significance was determined by Welch’s t-test. Mean ± S.D. is displayed. n.s., not significant; and \*\**p*<0.01. Scale bars: 50 μm.

**Supplemental Figure 5.**
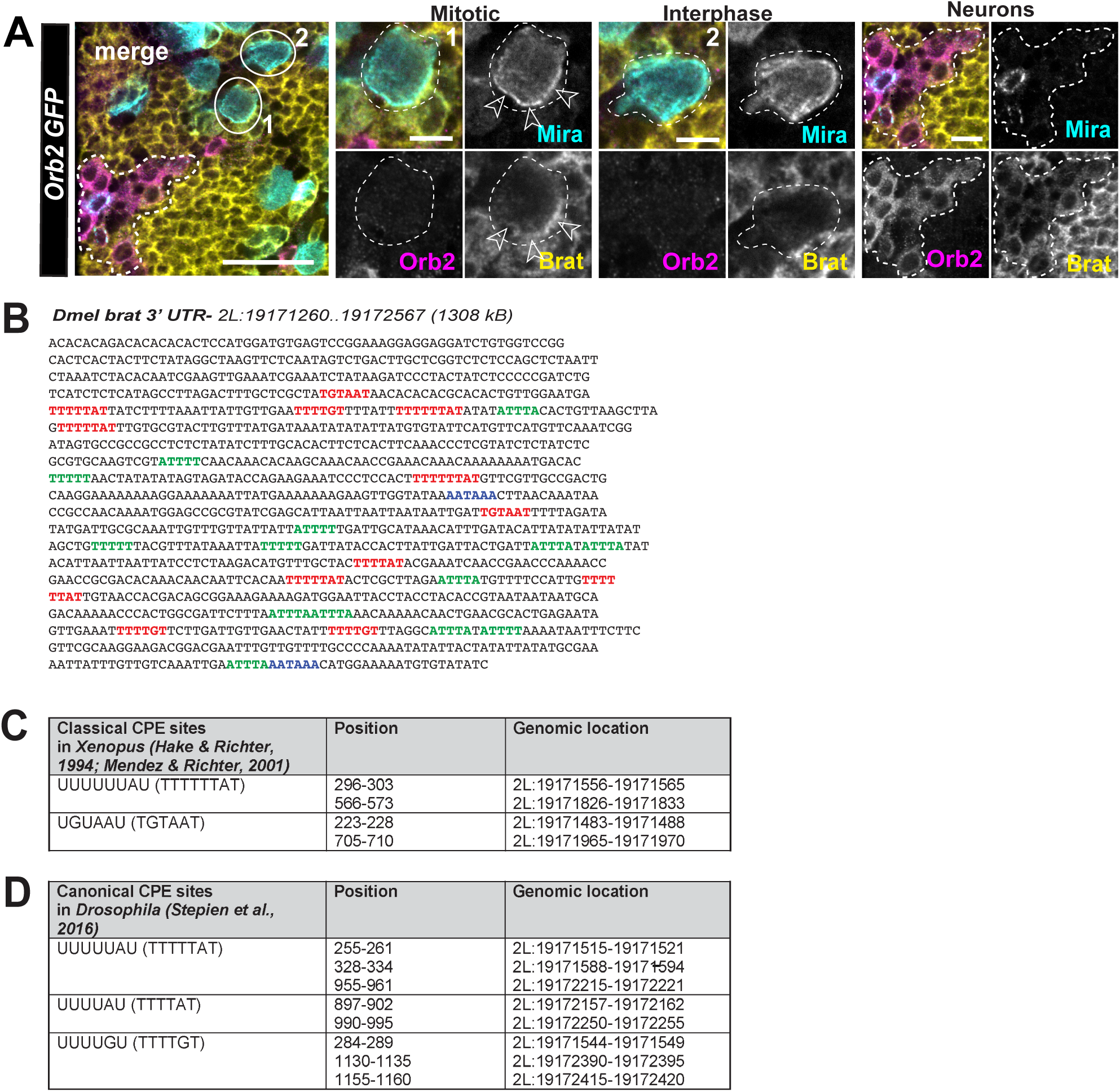
T**h**e ***brat* 3’UTR contains multiple CPEs.** (A) Orb2 distribution was examined in larval brains expressing*Orb2-GFP* and stained for GFP (Orb2, magenta), Brat (yellow), and Mira (cyan) antibodies. Insets show protein distributions within a mitotic NSC (circle 1), where Mira and Brat are localized to the basal crescent, and an interphase NSC (circle 2). The outlined area marks presumptive neurons positive for Orb2 and Brat; although, Orb2 and Brat appear to show mutually exclusive enrichments within these cells. (B) The *Drosophila melanogaster brat* 3’UTR sequence with cytoplasmic polyadenylation element (CPE; red), polyadenylation signal (PAS; blue), and AU-rich element (ARE; green) motifs annotated. (C) Genomic location of classical CPE sites (UUUUUUAU) and (UGUAAU), as identified by (Hake & Richter, 1994; Mendez & Richter, 2001) within the *brat* 3’UTR. (D) Genomic location of additional canonical CPE sites (UUUUUAU), (UUUUAU), and (UUUUGU), as identified by (Stepien et al., 2016) within the *brat* 3’UTR. Scale bars: 10 μm; insets, 5 μm.

